# FABP4 Couples Lipid Metabolism to PD-L1 Stabilization in Immunosuppressive Macrophages

**DOI:** 10.64898/2026.04.09.717546

**Authors:** Jianyu Yu, Jonathan Shilyansky, Jiaqing Hao, Anthony Avellino, Yanwen Sun, Xingshan Jiang, Zhaohua Wang, Xiaochun Han, Melissa A. Curry, Sonia L Sugg, Bing Li

## Abstract

Metabolic dysregulation in obesity reshapes immune function, but how lipid signals drive immune suppression remains unclear. Here, we identify a FABP4–PD-L1 axis that links lipid metabolism to immune checkpoint regulation in monocytes and macrophages. Single-cell transcriptomics revealed a distinct FABP4^high^ immunosuppressive macrophage subset enriched under high-fat diet (HFD) conditions, characterized by impaired antigen presentation and elevated PD-L1 expression. Mechanistically, palmitic acid (PA) induces FABP4 and promotes PD-L1 palmitoylation, leading to its stabilization on the cell surface independent of transcriptional regulation. FABP4 is essential for this process, which enables PD-L1 surface stabilization, immunosuppression and mammary tumor progression. In humans, a conserved CD14^int^CD16⁺ monocyte population exhibits elevated FABP4–PD-L1 signaling and correlates with obesity and invasive breast cancer. These findings establish PD-L1 as a metabolically regulated protein and reveal a mechanism by which lipid excess drives immune evasion, suggesting that targeting FABP4 may enhance responses to immune checkpoint blockade.

**Graphical Abstract:** 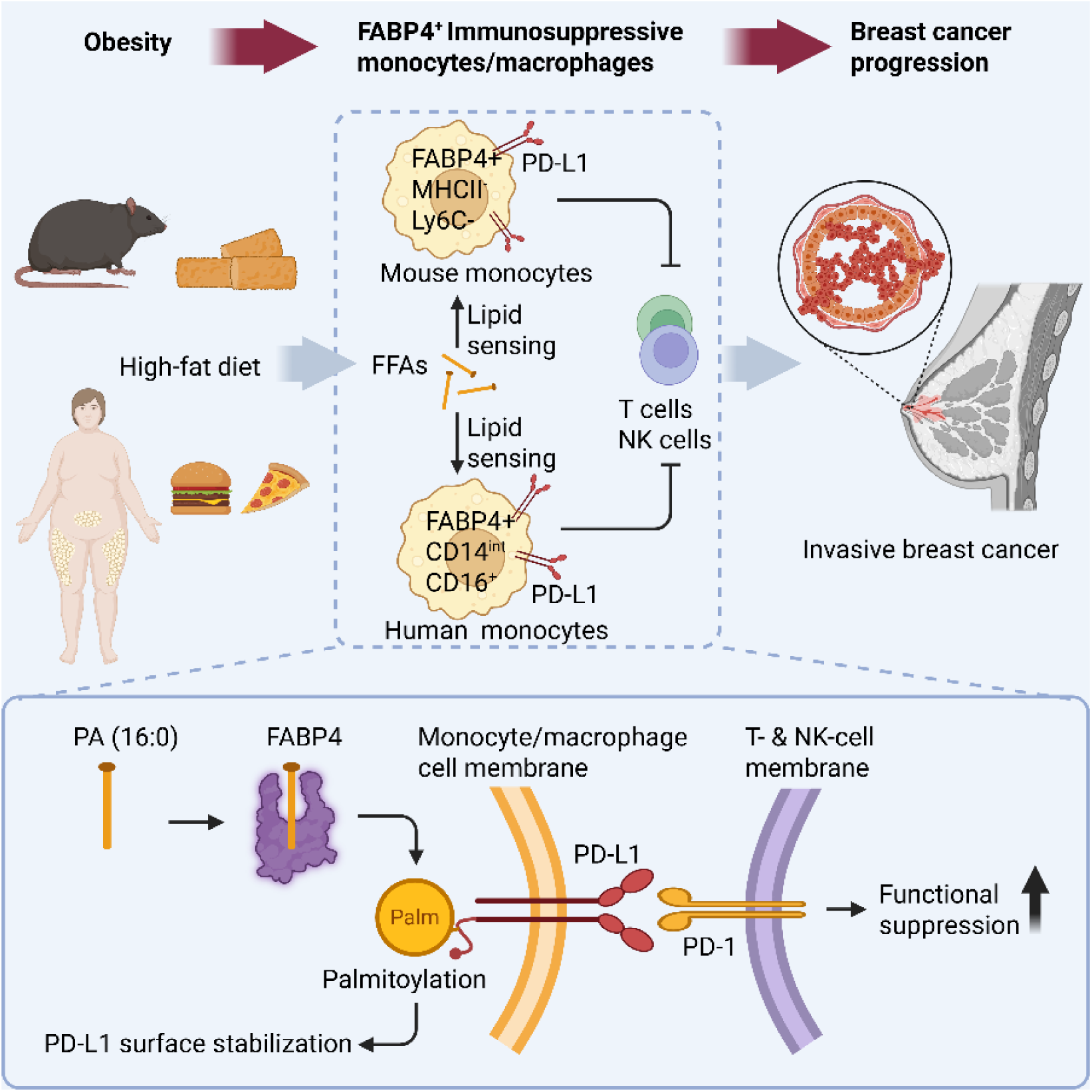

**Highlights:**

1. FABP4 defines a lipid-responsive, immunosuppressive monocyte/macrophage subset
2. FABP4 links lipid sensing to PD-L1 expression in macrophages
3. FABP4 enables palmitic acid-dependent PD-L1 palmitoylation and stabilization
4. FABP4–PD-L1 signaling correlates with obesity and invasive breast cancer in humans

## Introduction

Monocytes and macrophages are central components of the innate immune system, playing critical roles in tissue homeostasis, host defense, inflammation, or cancer^1–4^. However, their extensive heterogeneity has limited their utility as reliable diagnostic markers or therapeutic targets^5, 6^. For example, an elevated monocyte-to-lymphocyte ratio (MLR) has been associated with poor prognosis in breast cancer patients^7, 8^, yet its predictive power remains modest due to the functional diversity of circulating monocyte subsets^9^. Thus, defining the molecular mechanisms that specify distinct monocyte subsets and their functional states is essential for developing more robust prognostic tools and therapeutic strategies.

Metabolic reprograming has emerged as a key driver of monocyte heterogeneity across diverse disease contexts^10, 11^. In general, glycolytic metabolism is associated with pro-inflammatory and anti-tumoral functions, whereas lipid-oriented metabolic programs are linked to immunosuppressive and pro-tumoral phenotypes^12, 13^. This paradigm is particularly relevant in obesity-associated breast cancer, where systemic lipid dysregulation reshapes immune cell function^14, 15^. However, in the context of complex metabolic environments, such as obesity^16–18^, the mechanisms by which extrinsic lipid signals are integrated with intrinsic cellular programs to define monocyte fate and function remain poorly understood.

In breast cancer, circulating monocytes can express Programmed Cell Death Ligand 1 (PD-L1), a key immunosuppressive checkpoint molecule that suppresses T- and NK-cell activity and facilitates tumor progression and metastasis^19^. Although PD-1/PD-L1-targeted therapies have been explored clinically^20^, their efficacy in breast cancer remains limited, with low response rates to monotherapy^21, 22^. One major limitation is the persistence of myeloid-driven immunosuppression, including sustained induction of PD-L1 in monocytes through upstream signaling pathways that are not addressed by checkpoint blockade^23^. Therefore, identifying the mechanisms that regulate PD-L1 expression in monocytes is critical for improving immunotherapy outcomes. Emerging evidence further suggests that obesity-associated lipid dysregulation contributes to immunosuppressive programming in breast cancer^24, 25^. However, the specific monocyte subsets that sense lipid cues and the molecular mechanisms by which these signals drive PD-L1 expression have not been well defined.

Fatty acid-binding proteins (FABPs) are a family of intracellular lipid chaperones that facilitate the solubilization and trafficking of hydrophobic fatty acids across diverse tissues^26, 27^. Among them, adipocyte fatty acid-binding protein (A-FABP, also known as FABP4) is highly expressed in monocytes and macrophages and serves as key regulator of lipid metabolism and immune functions^28, 29^. In addition to its intracellular roles, FABP4 can be secreted as a circulating factor that promotes tumor progression^30–32^. Despite these diverse functions, whether FABP4 integrates systemic lipid signals with immunosuppressive programs in circulating monocytes, particularly in obesity-associated breast cancer, remains unclear.

In this study, we identified a distinct subset of FABP4^high^ monocytes and macrophages with potent immunosuppressive features using single-cell transcriptomics (scRNA-seq) and immune profiling. We show that these cells are highly responsive to dietary lipid signals and exhibit elevated PD-L1 expression and potent immunosuppressive functions. Mechanistically, FABP4 binds and responds to palmitic acids (PA) and promotes PD-L1 palmitoylation, leading to its stabilization on the cell surface. Translationally, we identify a conserved CD14^int^CD16^+^ monocyte subsets in human blood that exhibits elevated FABP4-PD-L1 signaling, correlated with obesity and associated with breast cancer invasiveness and systemic immune suppression. Collectively, our findings uncover a previously unrecognized FABP4-PD-L1 axis that links lipid dysregulation to immunosuppressive monocyte function in breast cancer progression.

## Results

### Single-cell transcriptomics identify an FABP4^+^ immunosuppressive macrophage subset in obesity

To characterize the role of immunosuppressive macrophages in obesity, C57BL/6 mice (n=3) were fed either a low-fat diet (LFD, 5% fat) or high-fat diet (HFD, 45% fat) for 3 months. F4/80^+^ splenic macrophages were isolated and subjected to single-cell RNA sequencing (scRNA-seq) (Figure 1A). A total of 17089 cells were profiled and classified into 13 distinct cell subsets based on canonical transcriptional markers (Figure 1B, and Figure S1A). Among these, we identified a distinct macrophage subset (macrophage_3) characterized by high expression of the lipid chaperone gene Fabp4 (Fabp4^hi^) (Figure 1C). Consistent with enhanced lipid handling functions^33^, Fabp4^hi^ macrophages exhibited upregulation of lipid metabolism-associated genes, such as Cd36 and Pparg (Figure 1D). These cells also showed increased expression of genes involved in cell-cell adhesion (e.g., L1cam and Itgal), T cell suppression (e.g., Bcl6 and Klf4), and TGFβ/SMAD signaling (e.g., Tgfbr1, Tgfbr2, Tgfbr3, Smad3, etc.), while downregulating genes related to antigen presentation and co-stimulation (e.g., MHCII components, Cd74, Cd80, Cd86, Flt3, etc.) (Figure 1D, S1B-S1F). Pathway enrichment analyses further supported a strongly immunosuppressive phenotype of Fabp4^hi^ macrophages with upregulation of cell interaction (Figure 1E) and downregulation of antigen-presentation and T cell activation pathways (Figure 1F). Additionally, Reactome pathway analysis showed enrichment of immunosuppressive features, including upregulation of TGFβ/SMAD signaling and CTLA inhibitory signaling (Figure S1G), alongside downregulation of cell division, CD28-related costimulatory signaling, and antigen presentation pathways (Figure S1H). Collectively, these data identify Fabp4^hi^ macrophages as lipid-enriched, immunosuppressive subsets with enhanced cell-cell interaction capacity but impaired antigen-presenting function.

**Figure 1.**
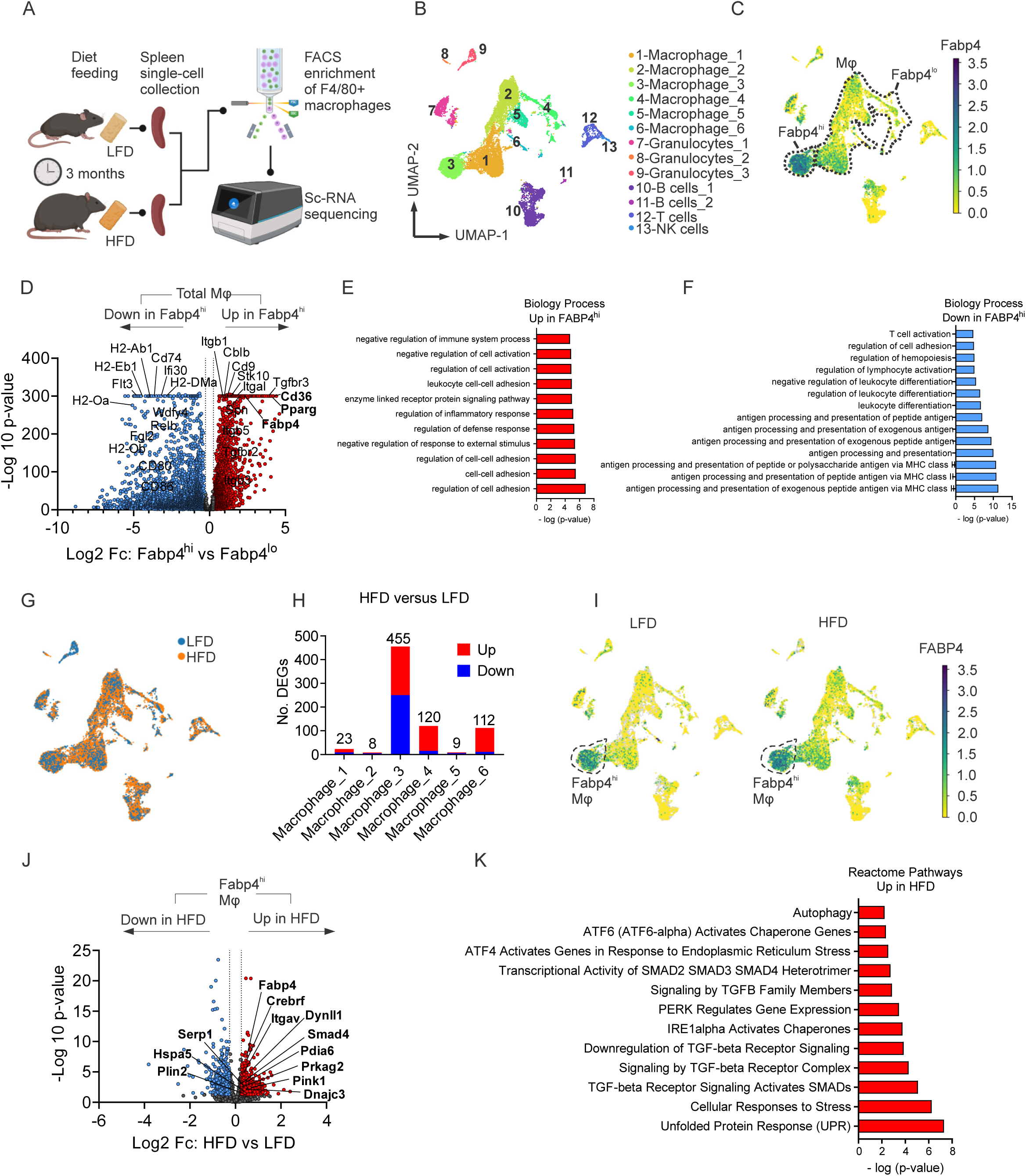
Single-cell transcriptomic analysis shows immune-suppressive function of Fabp4^hi^ macrophages in mouse spleen. (A) Schematic of single-cell RNA sequencing (ScRNA-seq) on FACS-enriched splenic F4/80^+^ macrophage from mice fed low-fat diet (LFD) or high-fat diet (HFD). (B) ScRNA-seq clustering analysis of F4/80^+^-enriched splenocytes from mice fed LFD (n=3) and HFD (n=3) with lineage annotation. (C) Identification of Fabp4^hi^ and Fabp4^lo^ splenic macrophage based on Fabp4 gene expression level in ScRNA-seq. Dash lines circle macrophages. (D) Differential expressing gene (DEG) analysis comparing Fabp4^hi^ and Fabp4^lo^ splenic macrophages shown in volcano plot with an adjusted p value cutoff of 0.05 and a Log2 fold change cutoff of ±0.26. Red dots are significantly upregulated genes in Fabp4^hi^, blue dots are significantly downregulated genes in Fabp4hi, and gray dots are genes with no significant difference. (E) Pathway enrichment analysis of significantly up-regulated genes in Fabp4^hi^ compared to Fabp4^lo^ splenic macrophages using Biology Process database. (F) Pathway enrichment analysis of significantly down-regulated genes in Fabp4^hi^ compared to Fabp4^lo^ splenic macrophages using Biology Process database. (G) Overlapping clustering distribution of splenocytes between mice fed HFD and LFD. (H) Amount of DEGs across different macrophage clusters in HFD versus LFD. (I) UMAP of Fabp4 gene expression in splenocytes from mice fed HFD and LFD. Dash lines circle Fabp4^hi^ splenic macrophages. (J). DEG analysis comparing Fabp4^hi^ splenic macrophages in HFD and LFD shown in volcano plot with an adjusted p value cutoff of 0.05 and a Log2 fold change cutoff of ±0.26. Red dots are significantly upregulated genes in HFD, blue dots are significantly downregulated genes in HFD, and gray dots are genes with no significant difference. (K) Pathway enrichment analysis of significantly upregulated genes in Fabp4^hi^ splenic macrophages from mice fed HFD compared to LFD using Reactome Pathways database. Also see Figure S1.

Given their lipid-associated features, we next examined the dietary regulation of Fabp4^hi^ macrophages (Figure 1G). Compared with other macrophage populations, Fabp4^hi^ macrophages (macrophage_3) exhibited the greatest transcriptional response to HFD (Figure 1H), including further induction of Fabp4 expression (Figure 1I, S1I-S1J). HFD also enriched pathways related to ER stress, autophagy, and TGFβ/SMAD signaling (Figure 1J, 1K), indicating that Fabp4^hi^ macrophages are highly responsive to metabolic stress and acquire enhanced immunosuppressive features under obesogenic conditions.

### FABP4^+^ monocytes/macrophages exhibit high PD-L1 expression and systemic immunosuppressive features in mice

To functionally characterize FABP4^+^ macrophages, we identified that Ly6c2 and H2-Eb1, macrophage differentiation-associated genes, were absent in the Fabp4^+^ subset (macrophage_3) (Figure 2A). Flow cytometric analysis stratified splenic macrophages into four subsets based on Ly6C and MHCII expression, which were Q1 (MHCII^-^Ly6C^+^), Q2 (MHCII^+^Ly6C^+^), Q3 (MHCII^+^Ly6C^-^), and Q4 (MHCII^-^Ly6C^-^) (Figure 2B, Figure S2A). Among these, FABP4 expression was predominantly enriched in the Q4 subset (Figure 2C).

**Figure 2.**
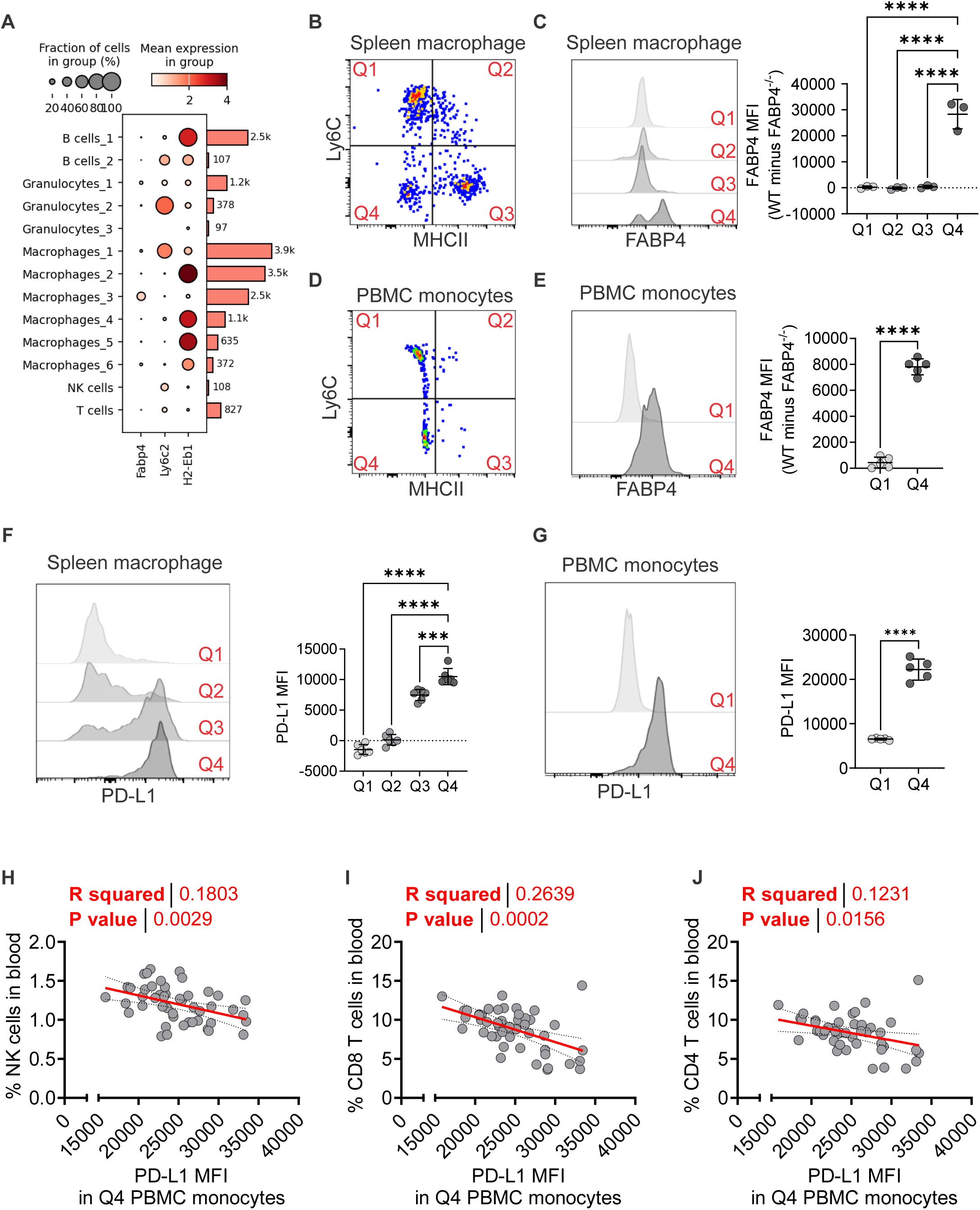
Flow cytometric analysis shows high PD-L1 expression in FABP4^hi^ monocyte in mice. (A) Dot plot of the relative expression of genes including Fabp4, Ly6c2, and H2-Eb1 across splenocyte subsets in scRNA-seq. (B) Splenic macrophage subsets (Q1-Q4) classification based on surface MHCII and Ly6C expression under gate CD11b^+^F4/80^+^ in flow cytometry. (C) Flow cytometric analysis of intracellular FABP4 levels across splenic macrophage subsets (Q1–Q4), shown as representative flow plots and quantified as relative intracellular FABP4 mean fluorescence intensity (MFI) by subtracting FABP4^-/-^ from WT. (D) PBMC monocyte subsets (Q1-Q4) classification based on surface MHCII and Ly6C expression under gate CD11b^+^F4/80^+^ in flow cytometry. (E) Flow cytometric analysis of intracellular FABP4 levels across PBMC monocyte subsets (Q1 and Q4), shown as representative flow plots and quantified as relative intracellular FABP4 MFI by subtracting FABP4^-/-^ from WT. (F) Flow cytometric analysis of surface PD-L1 levels in splenic macrophage subsets, shown as representative flow plots and quantified as PD-L1 MFI. (G) Flow cytometric analysis of surface PD-L1 levels in PBMC monocyte subsets, shown as representative flow plots and quantified as PD-L1 MFI. (H) Correlation analysis of surface PD-L1 MFI in Q4 PBMC monocyte subset and NK cell percentage in PBMC by flow cytometry. (I) Correlation analysis of surface PD-L1 MFI in Q4 PBMC monocyte subset versus CD8^+^ T cell percentage in PBMC by flow cytometry. (J) Correlation analysis of surface PD-L1MFI in Q4 PBMC monocyte subset versus CD4^+^ T cell percentage in PBMC by flow cytometry. Data are presented as the mean ± SD. *, *P* < 0.05; **, *P* < 0.01; ***, *P* < 0.001; ****, *P* < 0.0001; ns, nonsignificant; one-way ANOVA with Bonferroni’s multiple comparison test for (C) and (F), unpaired two-tailed t test for (E) and (G), and linear regression analysis for (H-J). Also see Figure S2.

Analysis of mouse peripheral blood mononuclear cells (PBMCs) confirmed that FABP4 expression was predominantly found in monocytes with low or absent expression in neutrophils and non-myeloid populations (Figure S2B, S2C). Although circulating monocytes were primarily distributed in the Q1 and Q4 compartments (Figure 2D, S2D), FABP4 expression was consistently enriched in the Q4 subset (Figure 2E), indicating a conserved FABP4^+^ Q4 subset (MHCII^-^Ly6C^-^) in both spleen and blood.

Given the immunosuppressive transcriptional profile of FABP4^+^ macrophages, we next examined PD-L1 expression. Strikingly, Q4 macrophages/monocytes exhibited the highest surface PD-L1 levels in both spleen and blood (Figure 2F, 2G). Importantly, PD-L1 expression in Q4, but not Q1, monocytes negatively correlated with circulating NK cells, CD8^+^ T cells, and CD4^+^ T cells (Figure 2H-2J, S2F-S2H), suggesting systemic immunosuppressive activity. Together, these findings identify FABP4^+^ Q4 monocytes as a dominant PD-L1^+^ immunosuppressive population under physiological conditions.

### HFD enhances the immunosuppressive function of FABP4^+^ monocytes in breast cancer models

Because FABP4 is closely linked to lipid sensing^34^, we next assessed whether HFD-induced obesity regulated FABP4^+^ monocyte function. PBMCs from mice fed LFD or HFD were characterized using flow cytometry (Figure 3A). HFD feeding increased FABP4 expression specifically in Q4 monocytes, but not the Q1 subset (Figure 3B), and expanded the proportion of Q4 monocytes in circulation (Figure 3C). Notably, HFD also selectively enhanced PD-L1 surface expression in Q4 monocytes (Figure 3D).

**Figure 3.**
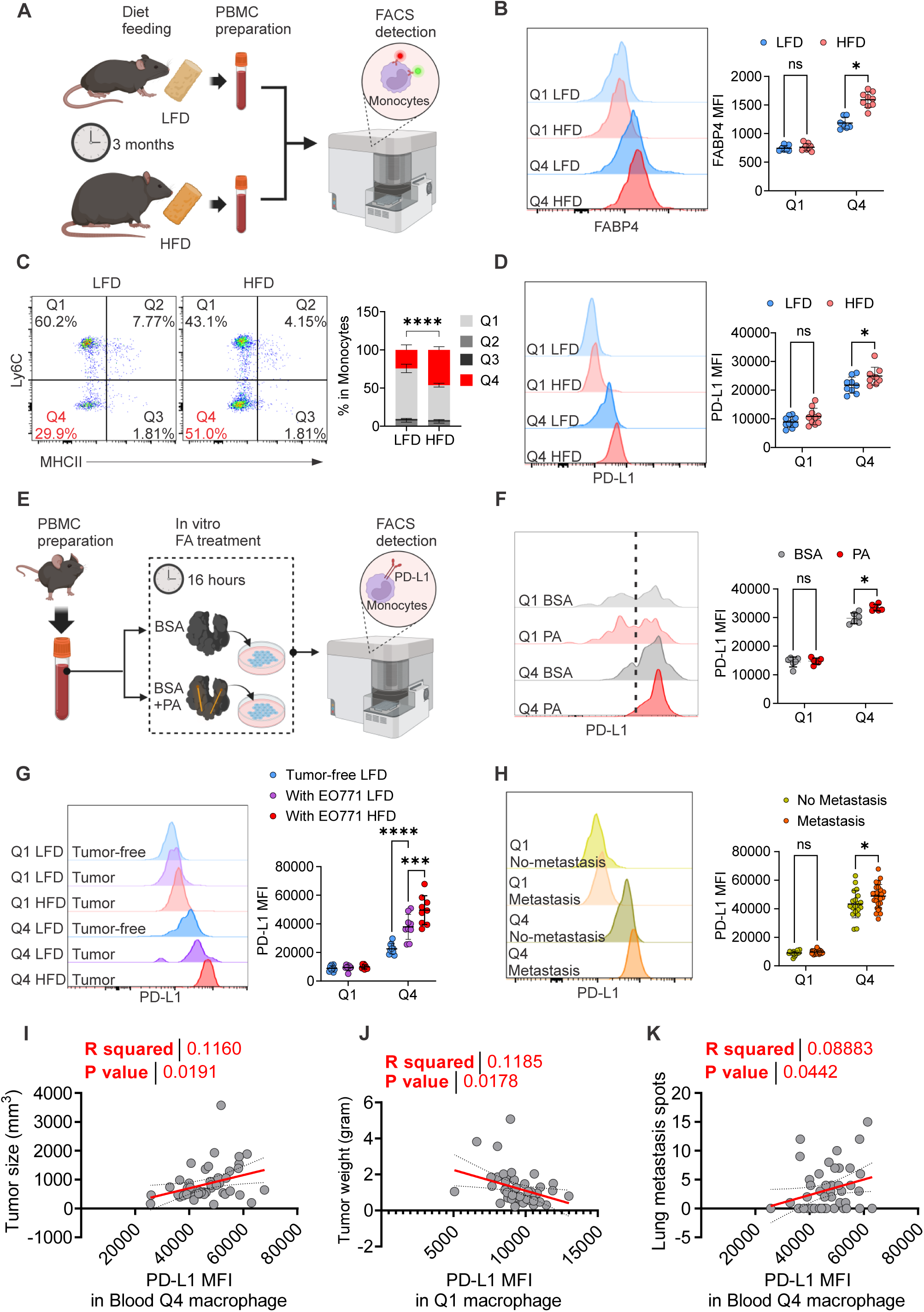
FABP4^+^ monocytes in mouse PBMC respond to lipid signals and contribute to breast cancer risk. (A) Schematic of flow cytometric analysis of PBMC monocytes from mice fed LFD or HFD for 3 months. (B) Flow cytometric analysis of intracellular FABP4 level in PBMC monocyte subsets (Q1 and Q4) in LFD and HFD, shown as representative flow plot and quantified intracellular FABP4 MFI. (C) Flow cytometric analysis of PBMC monocyte subset (Q1-Q4) compartments in LFD and HFD, shown as representative flow plot and quantified as ratios in monocytes. Statistical significance is indicated for the Q4 subset compartments between the two groups. (D) Flow cytometric analysis of surface PD-L1 levels in PBMC monocyte subsets (Q1 and Q4) from mice in LFD and HFD, shown as representative flow plot and quantified PD-L1 MFI. (E) Schematic of flow cytometric analysis of monocyte surface PD-L1 levels in PBMCs treated by BSA-conjugated palmitic acid (PA), and BSA-only control in vitro from 16 hours. (F) Flow cytometric analysis of surface PD-L1 levels in PBMC monocyte subsets (Q1 and Q4) treated with 200µM PAor BSA in vitro for 16 hours, shown as representative flow plot and quantified PD-L1 MFI. (G) Flow cytometric analysis of surface PD-L1 levels in PBMC monocyte subsets (Q1 and Q4) from mice fed LFD or HFD and received orthotopically injection of EO771 tumor cells into fourth mammary fat pad, shown as representative flow plot and quantified PD-L1 MFI. (H) Flow cytometric analysis of surface PD-L1 levels in PBMC monocyte subsets (Q1 and Q4) from mice with or without lung metastasis after orthotopically injection of EO771 tumor cells into fourth mammary fat pad, shown as representative flow plot and quantified PD-L1 MFI. (I) Correlation analysis of surface PD-L1 MFI by flow cytometry in Q4 PBMC monocyte subset versus endpoint tumor size after orthotopically injection of EO771 tumor cells into fourth mammary fat pad. (J) Correlation analysis of surface PD-L1 MFI by flow cytometry in Q4 PBMC monocyte subset versus endpoint tumor weight after orthotopically injection of EO771 tumor cells into fourth mammary fat pad. (K) Correlation analysis of surface PD-L1 MFI by flow cytometry in Q4 PBMC monocyte subset versus number of lung metastasis spots after orthotopically injection of EO771 tumor cells into fourth mammary fat pad. Data are presented as the mean ± SD. *, *P* < 0.05; **, *P* < 0.01; ***, *P* < 0.001; ****, *P* < 0.0001; ns, nonsignificant; two-tailed multiple t test for (B), (D), (F), and (H), two-way ANOVA with Tukey’s multiple comparison test for (G), unpaired two-tailed t-test for (C), and linear regression analysis for (I-K). Also see Figure S3.

To further determine the impact of dietary lipids, we treated PBMCs with either bovine serum albumin (BSA)-conjugated palmitic acids (PA) or BSA control *in vitro* (Figure 3E). PA significantly increased PD-L1 surface expression in Q4, but not Q1, monocytes (Figure 3F), indicating that lipid exposure directly promotes immunosuppressive activation of FABP4^+^ monocytes.

To assess functional relevance in cancer, we orthotopically implanted EO771 tumors into LFD- or HFD-fed mice. HFD significantly increased tumor growth (Figure S3A-S3C) and lung metastasis (Figure S3D). Tumor challenging further induced PD-L1 surface expression specifically in Q4 monocytes, with a more pronounced effect under HFD (Figure 3G). Importantly, PD-L1 levels in Q4 monocytes (Figure 3H-3K), but not Q1 monocytes (Figure S3E-S3G), positively correlated with tumor burden and lung metastatic nodules, and FABP4^+^ Q4 macrophages were enriched in tumor stroma under HFD (Figure S3H, S3I). Altogether, these results demonstrate that FABP4^+^ monocytes integrate dietary lipids and tumor signals to promote immunosuppressive, pro-tumor functions.

### FABP4 promotes PD-L1 stabilization through post-translational mechanisms

Although surface PD-L1 expression was markedly elevated in FABP4^+^ monocytes/macrophages, no differences were observed at the transcriptional level in these cells, as assessed among different macrophage populations in WT mice (Figure S4A) and the FABP4^+^ subsets between WT and FABP4^-/-^ mice (Figure S4B), suggesting that FABP4 modulates PD-L1 surface expression at a post-translational level.

To investigate this, we examined PD-L1 surface expression across multiple sources of monocytes/macrophages from WT or FABP4^-/-^ mice. Under non-polarized basal conditions, FABP4 deficiency consistently reduced surface PD-L1 in macrophage cell lines (Figure S4C, S4D, Figure 4A), primary peritoneal macrophages (Figure S4E, S4F) and PBMC monocytes (Figure S4G). Upon polarization to an immunosuppressive M2 phenotype, PD-L1 surface expression was upregulated in response to IL-4, but downregulated following IL-4 withdrawal in WT macrophages. In contrast, FABP4 deficiency abolished this dynamic regulation of PD-L1 (Figure 4B). Furthermore, inhibition of proteasomal degradation with MG132 significantly increased FABP4 levels (Figure 4C) and concomitantly stabilized surface PD-L1 (Figure 4D), supporting a role for intracellular FABP4 in promoting PD-L1 stability at the cell surface.

**Figure 4.**
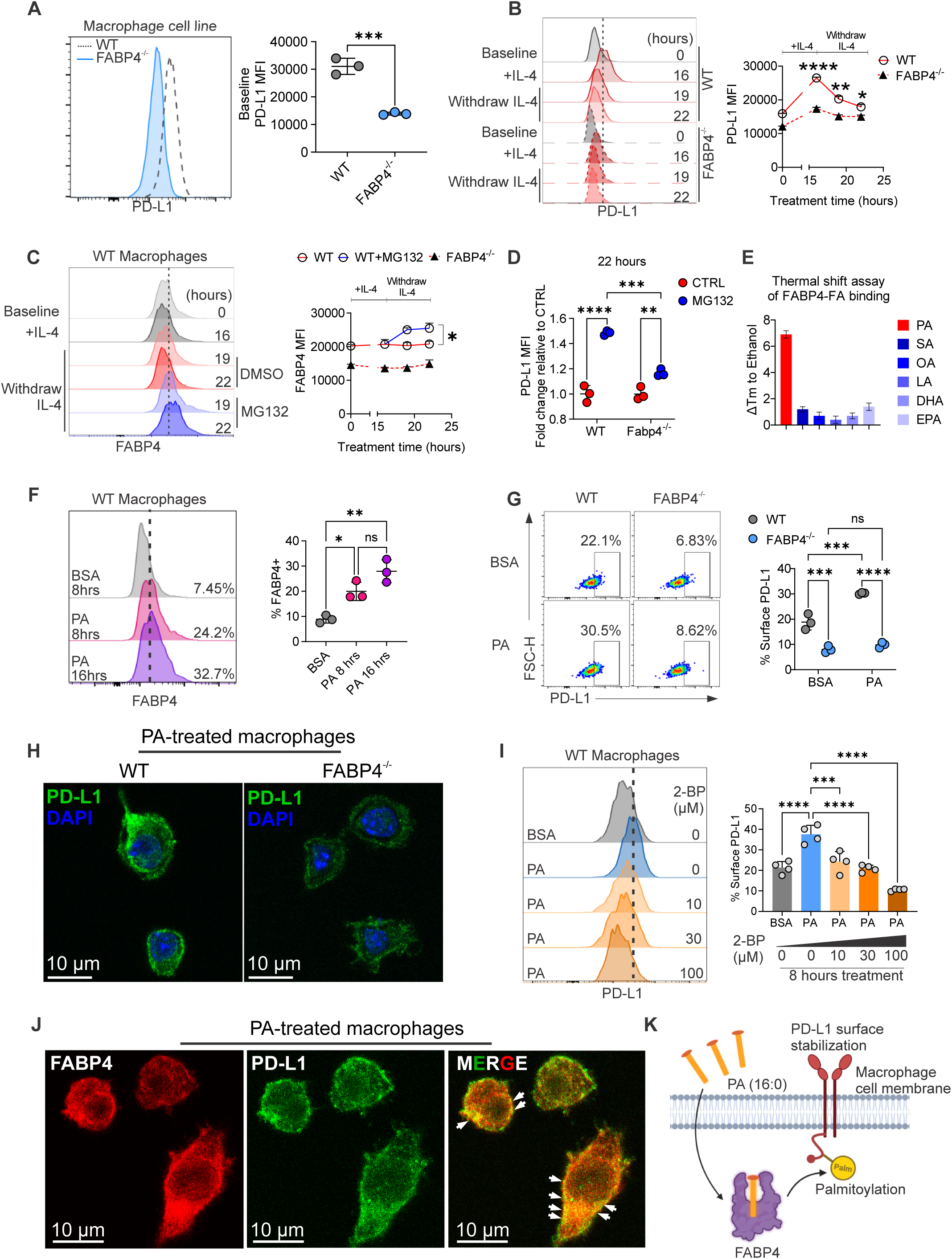
FABP4 senses lipids and stabilizes surface PD-L1 by mediating post-translational modification. (A) Baseline surface PD-L1 levels in WT and FABP4^-/-^ macrophages by flow cytometry, shown as representative flow plot and quantified surface PD-L1 MFI. (B) Surface PD-L1 kinetic in WT and FABP4^-/-^ macrophages across timepoints by flow cytometry, shown as representative flow plot and quantified surface PD-L1 MFI across timepoints. (C) Intracellular FABP4 kinetic in WT and FABP4^-/-^ macrophage across time in flow cytometry, shown as representative flow plot and quantified intracellular FABP4 MFI across timepoints. (D) Endpoint fold change of surface PD-L1 MFI by MG132 compared to DMSO control in WT and FABP4^-/-^ macrophages by flow cytometry. (E) Melting temperature shift (ΔTm) compared to ethanol control in thermal shift assays on purified FABP4 protein binding to different fatty acids (FA). (F) Flow cytometric analysis of intracellular FABP4 levels in WT macrophages treated with BSA or PA for 8 or 16 hours, shown as representative flow plot and quantified intracellular FABP4 MFI. (G) Surface PD-L1 levels in WT and FABP4^-/-^ macrophages treated with BSA or PA for 8 hours in vitro by flow cytometry, shown as representative flow plot and quantified surface PD-L1-positive percentages. (H) Measurement of PD-L1 expression in WT and FABP4^-/-^ macrophages by immunofluorescent staining. (I) Surface PD-L1 levels in WT macrophages treated BSA or PA in addition of different concentration of 2-bromopalmitate (2-BP) for 8 hours, shown as representative flow plot and quantified surface PD-L1-positive percentages. (J) Confocal staining of FABP4 (red) and PD-L1 (green) in WT macrophages treated PA for 8 hours. White arrows point to FABP4/PD-L1 colocalization areas (yellow). (K) Schematic of the mechanisms in which FABP4 mediates PA-induced PD-L1 palmitoylation and surface stabilization. Data are presented as the mean ± SD. *, *P* < 0.05; **, *P* < 0.01; ***, *P* < 0.001; ****, *P* < 0.0001; ns, nonsignificant; unpaired two-tailed t test for (A), one-way ANOVA with Bonferroni’s multiple comparison test for (F) and (I), two-way ANOVA with Tukey’s multiple comparison test for (B) (C), (D), and (G). Also see Figure S4.

Given that HFD increased PD-L1 expression in FABP4^+^ monocytes *in vivo* (Figure 3), we next sought to determine the mechanisms underlying FABP4-mediated lipid-induced regulation of surface PD-L1 expression. To identify the fatty acid ligand with the highest binding affinity for FABP4, we purified mouse FABP4 protein (Figure S4H) and performed thermal shift assays by incubating FABP4 protein with various dietary fatty acids. FABP4 showed the greatest melting temperature shift (ΔTm) upon incubation with PA compared with other fatty acids (Figure 4E), indicating the highest binding affinity. Additionally, PA treatment significantly increased FABP4 expression in macrophages (Figure 4F).

Based on these observations, we speculated that FABP4 mediates PA-induced PD-L1 surface expression. To test this, we first measured surface PD-L1 on WT and FABP4^-/-^ macrophages treated with or without PA. PA significantly increased surface PD-L1 levels, both in frequency and staining intensity, in WT macrophages, but not in FABP4^-/-^ macrophages (Figure 4G, S4I-S4J). This observation was further supported by immunofluorescence, which showed higher PA-induced PD-L1 surface intensity in WT than in FABP4^-/-^ macrophages (Figure 4H). As PA is known to enhance surface PD-L1 stabilization via palmitoylation^35, 36^, we inhibited palmitoylation using 2-bromopalmitate (2-BP). Notably, 2BP suppressed PA-induced surface PD-L1 upregulation in a dose-dependent manner on macrophages (Figure 4I). Confocal staining further demonstrated colocalization of FABP4 and PD-L1 near macrophage cell membrane following PA treatment (Figure 4J). Collectively, these findings support a model in which FABP4 facilitates PA-dependent PD-L1 palmitoylation, thereby stabilizing PD-L1 on the macrophage surface (Figure 4K).

### A conserved CD14^int^CD16^+^ monocyte subset exhibits immune-suppressive features in humans

Given the phenotypic and functional heterogeneity of monocytes/macrophages identified in mice, we next analyzed immune profiling of peripheral blood mononuclear cells (PBMCs) from non-cancer donors (Figure 5A and S5A). Under basal conditions, human PBMC monocytes were classified with two major populations: CD14^+^CD16^-^ and CD14^int^CD16^+^ subsets (Figure 5B). Notably, surface PD-L1 was highly expressed on CD14^int^CD16^+^ compared with CD14^+^CD16^-^ monocytes (Figure 5C). Correlation analysis showed a positive correlation of PD-L1 expression in CD14^int^CD16^+^, but not in CD14^+^CD16^-^ monocytes, with NK cell death (Figure 5D, S5B), suggesting an immune-suppressive activity of CD14^int^CD16^+^ monocytes in humans.

**Figure 5.**
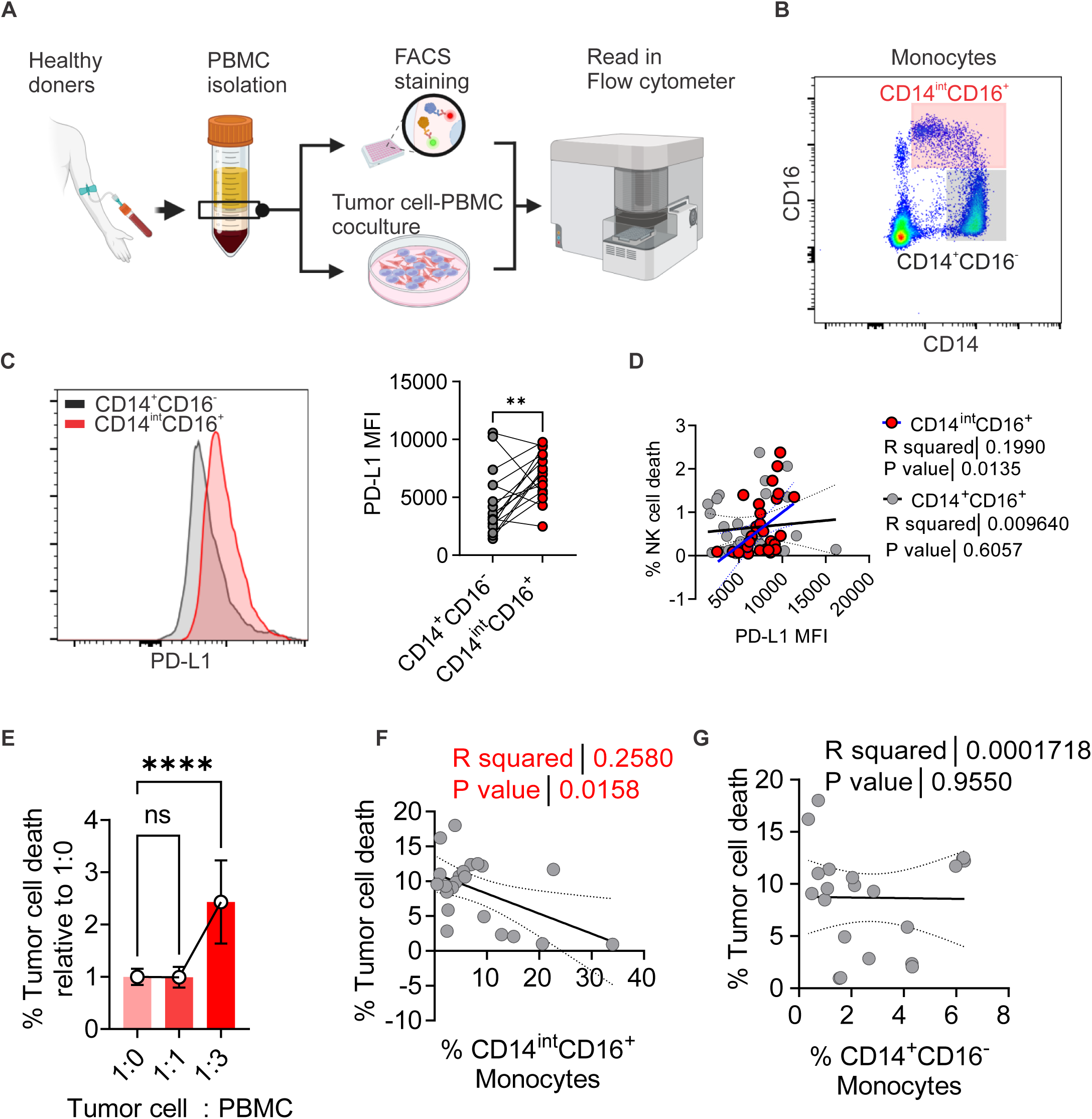
Conserved immunosuppressive CD14^int^CD16^+^ monocytes in human blood. (A) Schematic of PBMC isolation from non-cancer doners for immune profiling and coculture assay with tumor cells using flow cytometry. (B) Human PBMC monocyte classification based on CD16 and CD14 expression under gate CD56^-^ in flow cytometry. (C) Surface PD-L1 levels in human PBMC CD14^+^CD16^-^ and CD14^int^CD16^+^ monocytes in flow cytometry, shown as representative flow plot and quantified surface PD-L1 MFI. (D) Correlation analysis of surface PD-L1 MFI in CD14^+^CD16^-^ and CD14^int^CD16^+^ monocytes versus percentage of NK cell death in human PBMC. (E) Percentage of dead MDA cells in coculture with different ratios of PBMCs in vitro by flow cytometry. (F) Correlation analysis of CD14^int^CD16^+^ monocyte ratio versus percentage of MDA tumor cell death in coculture under tumor cell/PBMC ratio of 1:3. (G) Correlation analysis of CD14^+^CD16^-^ monocyte ratio versus percentage of MDA tumor cell death in coculture under tumor cell/PBMC ratio of 1:3. Data are presented as the mean ± SD. *, *P* < 0.05; **, *P* < 0.01; ****, *P* < 0.0001; ns, nonsignificant; paired two-tailed t test for (C), linear regression analysis for (D), (F) and (G), one-way ANOVA with Bonferroni’s multiple comparison test for (E). Also see Figure S5.

To further determine the function of CD14^int^CD16^+^ monocytes when exposed to tumor cells, we cocultured human PBMCs with breast cancer cell line, MDA-MB-231, at different cancer cell-PBMC ratios, and measured cancer cell death using flow cytometry (Figure S5C). CD45^-^ cancer cell death was increased when the ratio of human PBMCs elevated in the coculture system (Figure 5E), suggesting that PBMCs exert cytotoxic effects on tumor cells. Notably, the frequency of CD14^int^CD16^+^ monocytes (Figure 5F), but not CD14^+^CD16^-^ monocytes (Figure 5G), negatively correlated with tumor cell death (Figure 5F-5G), and abundance of both CD4^+^ and CD8^+^ T cells (Figure S5D and S5E). These results suggest a conserved CD14^int^CD16^+^ monocyte subsets exhibiting a pro-tumor immunosuppressive role in human blood.

### Obesity-related CD14^int^CD16^+^ monocytes correlate with invasive breast cancer

Given the immunosuppressive role of CD14^int^CD16^+^ monocytes in non-cancer donors, we next investigated the role of this monocyte subset in patients diagnosed with breast cancer. We collected PBMCs from patients with invasive or non-invasive breast cancer (Figure 6A, Table S1) and analyzed their immune profiling using the indicated flow cytometry gating strategy (Figure S5A). PBMCs in these patients mainly included T cells (60%), monocytes (10%), NK cells (10%), B cells (10%), and γδ T cells (<1%) (Figure S6A). To determine the potential roles of individual immune subsets in breast cancer progression, we found that invasive disease was associated with increased CD14^int^CD16^+^ monocytes, and reduced CD8^+^ T cells compared with non-invasive disease (Figure 6B-6C). No differences were found in other immune cells (Figure 6D-6H).

**Figure 6.**
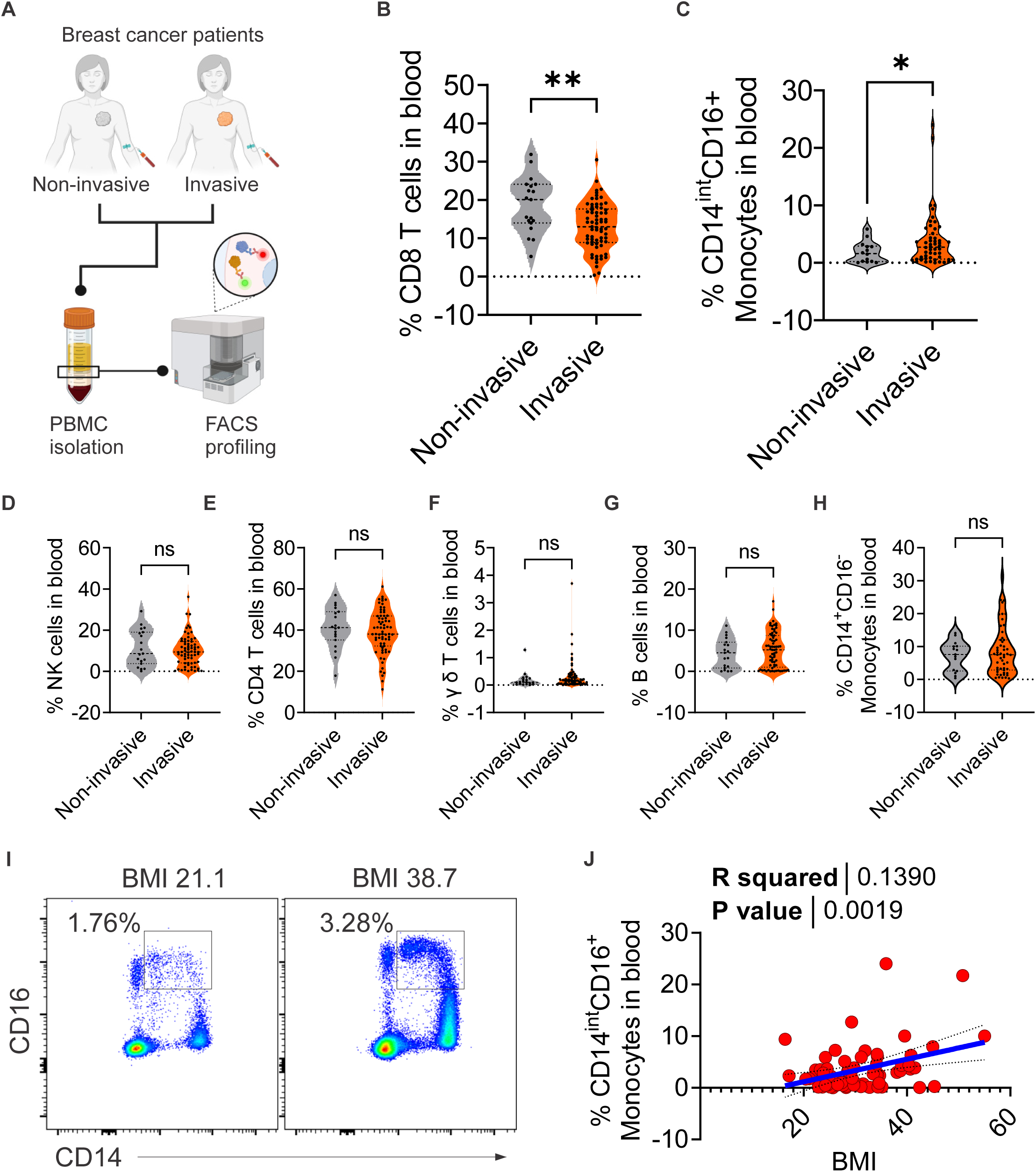
Human CD14^int^CD16^+^ monocytes mark systemic immunosuppression, obesity, and breast cancer invasiveness. (A) Schematic of PBMC isolation from clinical patients with non-invasive or invasive breast cancer types for immune profiling using flow cytometry. (B) Percentage of CD8^+^ T cells in PBMCs from patients with non-invasive and invasive breast cancer by flow cytometry. (C) Percentage of CD14^int^CD16^+^ monocytes in PBMCs from patients with non-invasive and invasive breast cancer by flow cytometry. (D) Percentage of NK cells in PBMCs from patients with non-invasive and invasive breast cancer by flow cytometry. (E) Percentage of CD4^+^ T cells in PBMCs from patients with non-invasive and invasive breast cancer by flow cytometry. (F) Percentage of γ δ T cells in PBMCs from patients with non-invasive and invasive breast cancer by flow cytometry. (G) Percentage of B cells in PBMCs from patients with non-invasive and invasive breast cancer by flow cytometry. (H) Percentage of CD14^+^CD16^-^ monocytes in PBMCs from patients with non-invasive and invasive breast cancer by flow cytometry. (I) Representative flow plots of CD14^int^CD16^+^ monocyte ratio in PBMCs from patients with lower and higher BMI in flow cytometry. (J) Correlation analysis of patient BMI versus CD14^int^CD16^+^ monocyte ratio in PBMCs. Data are presented as the mean ± SD. *, *P* < 0.05; **, *P* < 0.01; ****, *P* < 0.0001; ns, nonsignificant; unpaired two-tailed t-test for (B-H), linear regression analysis for (J). Also see Figure S6, Table S1.

Given that obesity increases breast cancer risk and progression^17, 18^, we next investigated whether obesity enhanced immunosuppressive monocytes in this cohort. Notably, patient BMI positively correlated with CD14^int^CD16^+^ monocyte frequency (Figure 6I-6J), but not with other immune cell types (Figure S6B), linking obesity to expansion of this immunosuppressive monocyte subset. Further correlation analyses showed that CD8^+^ T cells negatively correlated with age of patients (Figure S6C), NK cells negatively correlated with higher tumor stages (Figure S6D) and grades (Figure S6E). No significant differences were found among other immune cells and clinical parameters (Figure S6F). These findings demonstrate a consistent immunosuppressive CD14^int^CD16^+^ monocyte subset, which links obesity-associated breast cancer progression.

### FABP4-PD-L1 signaling defines immunosuppressive CD14^int^CD16^+^ monocytes in human breast cancer

Given the critical role of FABP4-PD-L1 signaling in promoting mammary tumor development in mouse models, we further investigated this signaling in human breast cancer. Consistent with non-cancer cohorts, surface PD-L1 levels were highly expressed in CD14^int^CD16^+^ monocytes (Figure S7A). Interestingly, surface PD-L1 was selectively elevated in CD14^int^CD16^+^ monocytes (Figure 7A), but not in CD14^+^CD16^-^ monocytes (Figure S7B), in invasive breast cancer. Furthermore, FABP4 expression was also enriched in this subset (Figure 7B), and selectively upregulated in invasive breast cancer. (Figure 7C, Figure S7C). Importantly, FABP4 expression positively correlated with surface PD-L1 expression in this subset (Figure 7D), indicating a conserved FABP4-PD-L1 axis. To further determine the immunosuppressive function of CD14^int^CD16^+^ monocytes, we assessed cytokine production in T cells and NK cells. Surface PD-L1 expression in CD14^int^CD16^+^ monocytes inversely correlated with TNFα production in NK cells and IL-2 production in NK cells, CD8 T cells, and CD4 T cells (Figure 7E-7H), supporting suppression of anti-tumor immunity in patients with breast cancer. Further analyses showed no correlation between either FABP4 or surface PD-L1 levels with other clinical parameters, including histology grading, menopausal status, BMI, or age (Figure S7D-S7G). Altogether, these findings establish FABP4-PD-L1 signaling in circulating CD14^int^CD16^+^ monocytes as a conserved immunosuppressive axis linking obesity to breast cancer progression.

**Figure 7.**
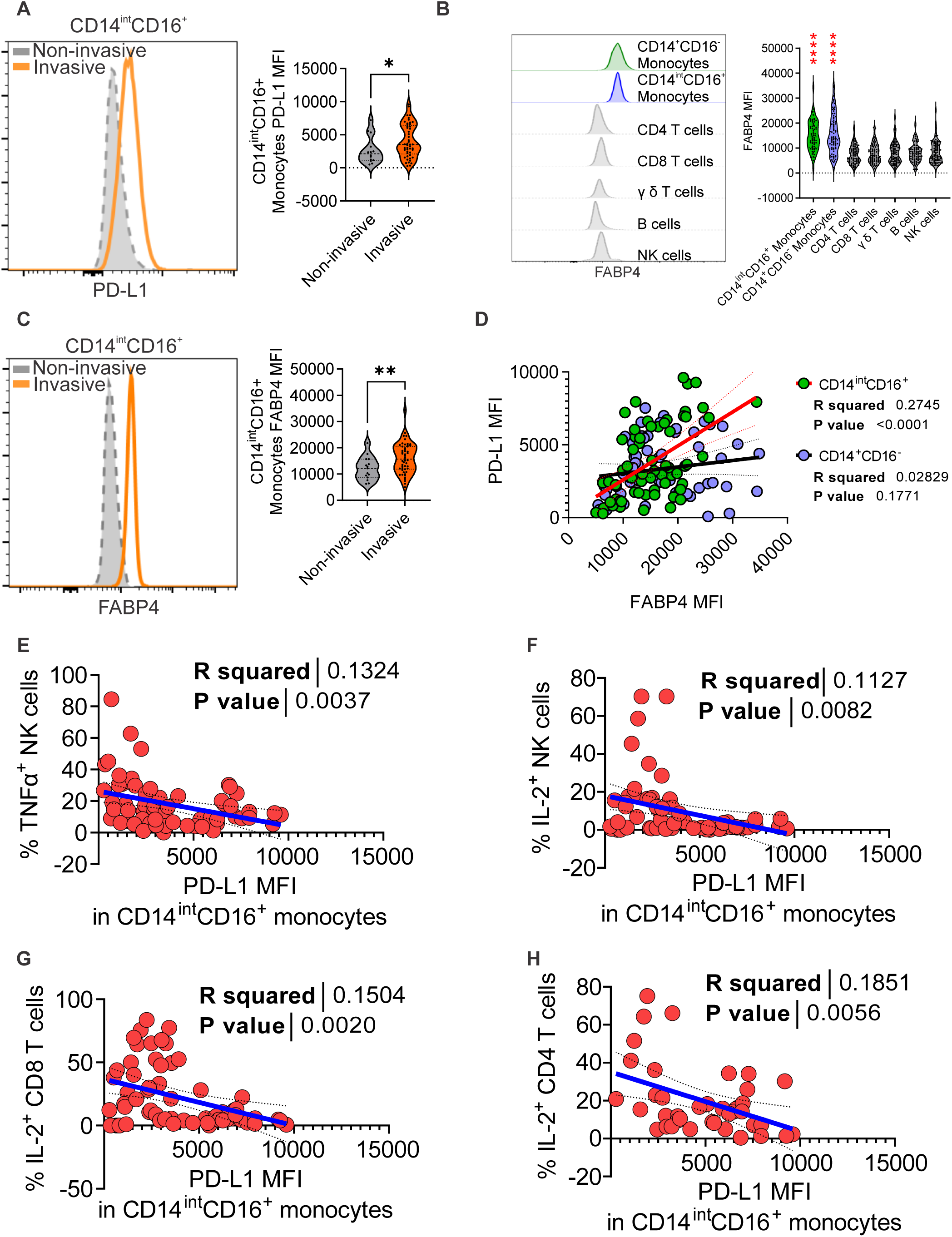
FABP4-PD-L1 signaling in human monocytes associates with systemic immunosuppression in breast cancer. (A) Surface PD-L1 levels in CD14^int^CD16^+^ monocytes from patients with non-invasive and invasive breast cancer types by flow cytometry, shown as representative flow plot and quantified surface PD-L1 MFI. (B) Intracellular FABP4 levels in different PBMC subsets in breast cancer patients by flow cytometry, shown as shown as representative flow plot and quantified intracellular FABP4 MFI. (C) Intracellular FABP4 levels in CD14^int^CD16^+^ monocytes from patients with non-invasive and invasive breast cancer types by flow cytometry, shown as representative flow plot and quantified intracellular FABP4 MFI. (D) Correlation analysis of intracellular FABP4 MFI versus surface PD-L1 MFI in CD14^int^CD16^+^ and CD14^+^CD16^-^ monocytes in breast cancer patients. (E) Correlation analysis of PD-L1 MFI in CD14^int^CD16^+^ monocytes versus percentage of TNFα^+^ in NK cells in PBMC from breast cancer patients. (F) Correlation analysis of PD-L1 MFI in CD14^int^CD16^+^ monocytes versus percentage of IL-2^+^ in NK cells in PBMC from breast cancer patients. (G) Correlation analysis of PD-L1 MFI in CD14^int^CD16^+^ monocytes versus percentage of IL-2^+^ in CD8^+^ T cells in PBMC from breast cancer patients. (H) Correlation analysis of PD-L1 MFI in CD14^int^CD16^+^ monocytes versus percentage of IL-2^+^ in CD4^+^ T cells in PBMC from breast cancer patients. Data are presented as the mean ± SD. *, *P* < 0.05; **, *P* < 0.01; ****, *P* < 0.0001; ns, nonsignificant; unpaired two-tailed t-test for (A) and (C), one-way ANOVA with Bonferroni’s multiple comparison test for (B). linear regression analysis for (D-H) Also see Figure S7.

## Discussion

A central finding of this study is the identification of FABP4 in regulation immunosuppression through promoting PD-L1 surface stability in monocytes/macrophages. While PD-L1 is classically regulated at the transcriptional level by inflammatory signaling pathways^37^, our data demonstrate a distinct mechanism in which lipid metabolism directly governs PD-L1 stability at the post-translational level. Despite minimal changes in PD-L1 mRNA, FABP4 deficiency markedly reduced PD-L1 surface expression, indicating that FABP4 regulates PD-L1 independently of transcription.

Mechanistically, we demonstrate that PA-induced PD-L1 surface expression requires FABP4 and depends on palmitoylation, a lipid modification known to prevent PD-L1 ubiquitination and lysosomal degradation^35, 38^. Inhibition of palmitoylation abolished PA-induced PD-L1 surface accumulation, supporting a model in which FABP4 facilitates PA-dependent palmitoylation to stabilize PD-L1 at the plasma membrane. In this framework, FABP4 functions as a lipid chaperone that enables substrate availability and/or spatial delivery of fatty acids for PD-L1 modification, effectively converting PD-L1 from a rapidly turned-over protein into a stabilized immune checkpoint receptor. More broadly, these findings identify PD-L1 as a metabolically regulated protein whose stability is directly controlled by lipid availability, establishing a previously unrecognized mechanism by which metabolic lipid flux shapes immune checkpoint dynamics.

This mechanism provides a conceptual advance in understanding how metabolic environments shape immune cell function^39, 40^. Rather than passively reflecting obesity, lipid excess actively instructs immune suppression^41^. In this context, FABP4 emerges as a central mediator linking extracellular lipid cues to immune checkpoint regulation. Our data demonstrate that dietary PA binds FABP4 with high-affinity and induces FABP4 expression in macrophages, likely through reported mechanisms involving ER stress and/or PPARγ signaling^42–44^. These findings extend the functional repertoire of FABP4 beyond lipid transport and energy metabolism, positioning it as an active regulator of immunosuppressive signaling in macrophages.

Importantly, the FABP4-PD-L1 axis integrated with broader FABP4-dependent programs that collectively reinforce a pro-tumor macrophage state. Prior studies have demonstrated that FABP4 promotes IL-6 production and STAT3 activation in macrophages, enhancing tumor growth and immune evasion^28^. In parallel, FABP4 has been shown to drive lipid droplet accumulation in macrophages, a metabolic phenotype associated with tumor metastasis^34^. Together with our current findings, these observations support a unified model in which FABP4 orchestrates a multidimensional pro-tumor macrophage program. Specifically, FABP4 coordinates: (1) cytokine signaling (IL-6/STAT3), (2) metabolic reprogramming (lipid droplet formation), and (3) immune checkpoint stabilization (PD-L1 palmitoylation). These processes are likely interconnected; for example, lipid droplets may serve as reservoirs of fatty acids that sustain palmitoylation reactions, while IL-6/STAT3 signaling has been reported to further enhance PD-L1 expression^45^. Thus, FABP4 functions as a central metabolic node that integrates lipid handling with immune regulatory circuits.

Our study also provides a framework for understanding monocytes and macrophage heterogeneity from a metabolic perspective. Using single-cell transcriptomics, we identify a FABP4^high^ immunosuppressive subset that is highly responsive to dietary lipid signals and characterized by coordinated suppression of antigen presentation alongside enhanced immune regulatory pathways. These findings support the concept that metabolic state is a primary determinant of monocyte/macrophage identity, particularly in metabolically perturbed conditions such as obesity and cancer^46^. The relevance of the FABP4–PD-L1 axis is further supported in human disease. We identify a CD14^int^CD16⁺ monocyte population that exhibits elevated FABP4 and PD-L1 expression, displays immunosuppressive features, and is expanded in obesity and patients with invasive breast cancer. These findings suggest that systemic metabolic dysregulation can precondition circulating monocytes toward a pro-tumor phenotype, potentially contributing to immune evasion prior to tumor infiltration.

Notably, our findings also provide mechanistic insight into the emerging “obesity paradox” observed in cancer immunotherapy, where obese patients exhibit improved responses to PD-1/PD-L1 blockade despite having worse baseline cancer risk^47^. One proposed explanation is that obesity induces a heightened PD-1/PD-L1–dependent immunosuppressive state, rendering tumors more susceptible to checkpoint blockade^48^. Our data support and extend this model by identifying FABP4 as a key metabolic driver of PD-L1 stabilization in macrophages under lipid-rich conditions. In this context, obesity-associated lipid excess, particularly saturated fatty acids such as PA, enhances FABP4 expression, leading to increased PD-L1 stabilization and immune suppression, thereby creating a checkpoint-dependent immunosuppressive state that is more responsive to PD-1/PD-L1 blockade. These findings suggest that FABP4-mediated lipid signaling may represent a critical upstream determinant of immunotherapy responsiveness and could help stratify patients or guide combination therapies.

From a therapeutic perspective, these results highlight an important opportunity. While current immune checkpoint inhibitors target PD-1/PD-L1 interactions, they do not address the upstream metabolic processes that regulate PD-L1 stability^49^. Targeting FABP4 or lipid-dependent post-translational modification pathways may therefore provide a strategy to disrupt immune checkpoint expression at its metabolic source, potentially enhancing immunotherapy efficacy and overcoming resistance driven by myeloid cells.

In summary, we define a FABP4–PD-L1 axis as a mechanistic link between lipid metabolism and immune checkpoint regulation in monocytes/macrophages. By coupling lipid sensing to PD-L1 palmitoylation and stabilization, FABP4 enables metabolic control of immune suppression. These findings establish a conceptual framework in which nutrient availability directly governs immune checkpoint dynamics through lipid-dependent protein modification and identify FABP4 as a key regulator of this process in obesity-associated breast cancer.

## Limitations

Several questions remain regarding the precise mechanisms underlying the FABP4-PD-L1 axis. While our data demonstrate a requirement for FABP4 in PA-induced PD-L1 palmitoylation, it remains unclear whether FABP4 directly regulates activity of palmitoyltransferases. In particular, the identity of the DHHC family palmitoyltransferase(s) mediating PD-L1 palmitoylation^36, 50^ in macrophages and their relationship to FABP4-dependent lipid trafficking remains to be defined. Additionally, other fatty acids may enhance PD-L1 surface expression, suggesting that distinct lipid species may differentially regulate immune checkpoint dynamics through overlapping or parallel mechanisms. Finally, while our human data establish clinical relevance, further studies will be required to determine whether targeting FABP4 can enhance responses to immune checkpoint blockade in patients.

## Acknowledgments

We thank the Department of Pathology at the University of Iowa for the access to the Cytek Aurora flow cytometer. B.L. thanks the funding support from NIH grants R01AI137324, R01CA180986, and U01CA272424 (part of National Cancer Institute Metabolic Dysregulation and Obesity Cancer Risk Program).

## Author contributions

J.Y., J.S., J.H., A.A., Y.S. J.X., W.Z., performed experiments and analyzed the data. X.H. maintained lean and obese mouse models, M.C, S.S., collected patient specimens and helped with clinical parameter analysis. B.L. designed the experiments. S.S., provided intellectual input and helped with paper writing. J.Y., and B.L. wrote the paper.

## Star Methods

### RESOURCE AVAILABILITY

#### Lead contact

Further information and requests for reagents may be directed to and will be fulfilled by the Lead Contact, Bing Li.

#### Material availability

All unique/stable reagents generated in this study are available from the lead contact with a completed materials transfer agreement.

#### Data and code availability

All data generated in this study will be made available upon request from the lead contact. This paper does not report original code. Any additional information required to reanalyze the data reported in this paper is available from the lead contact upon request.

## KEY RESOURCE TABLE

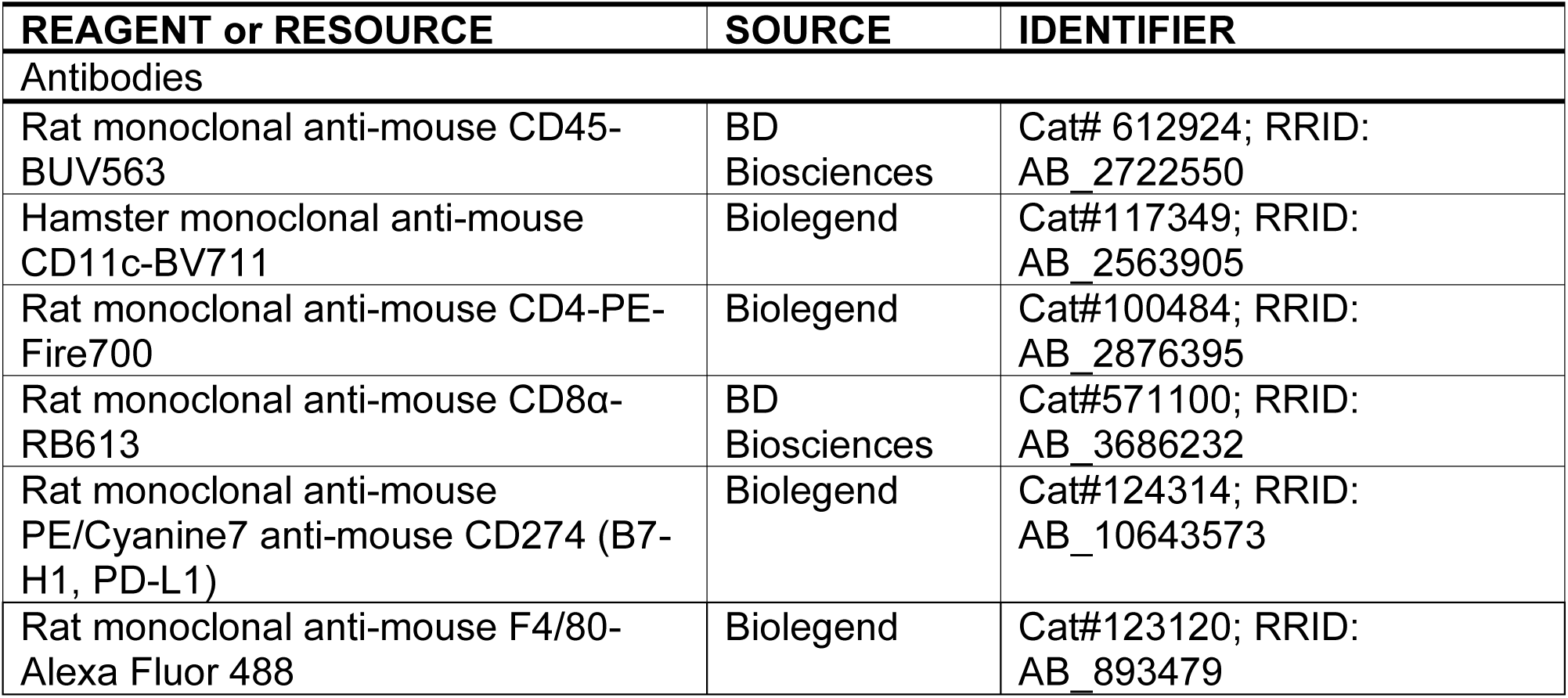
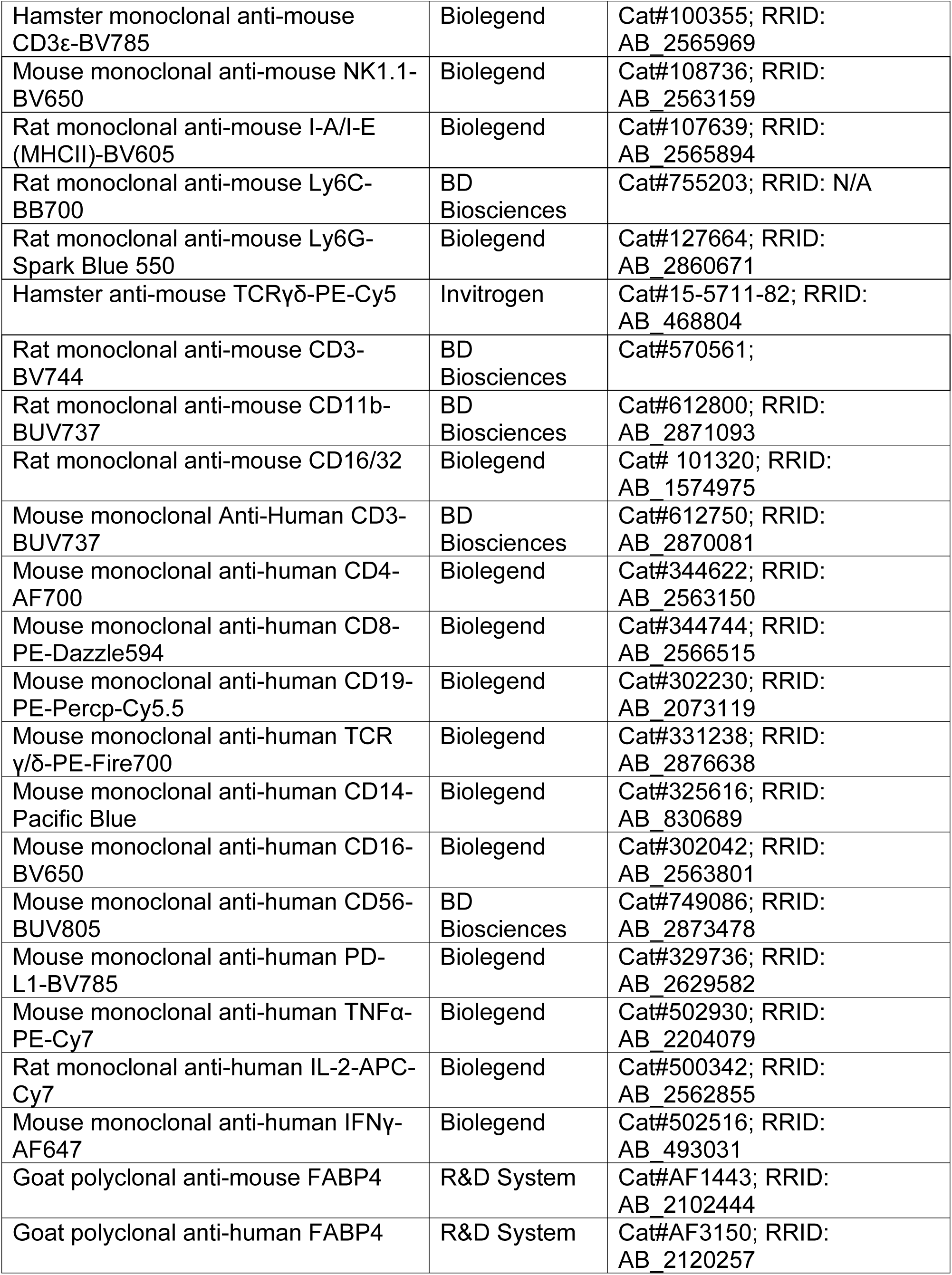
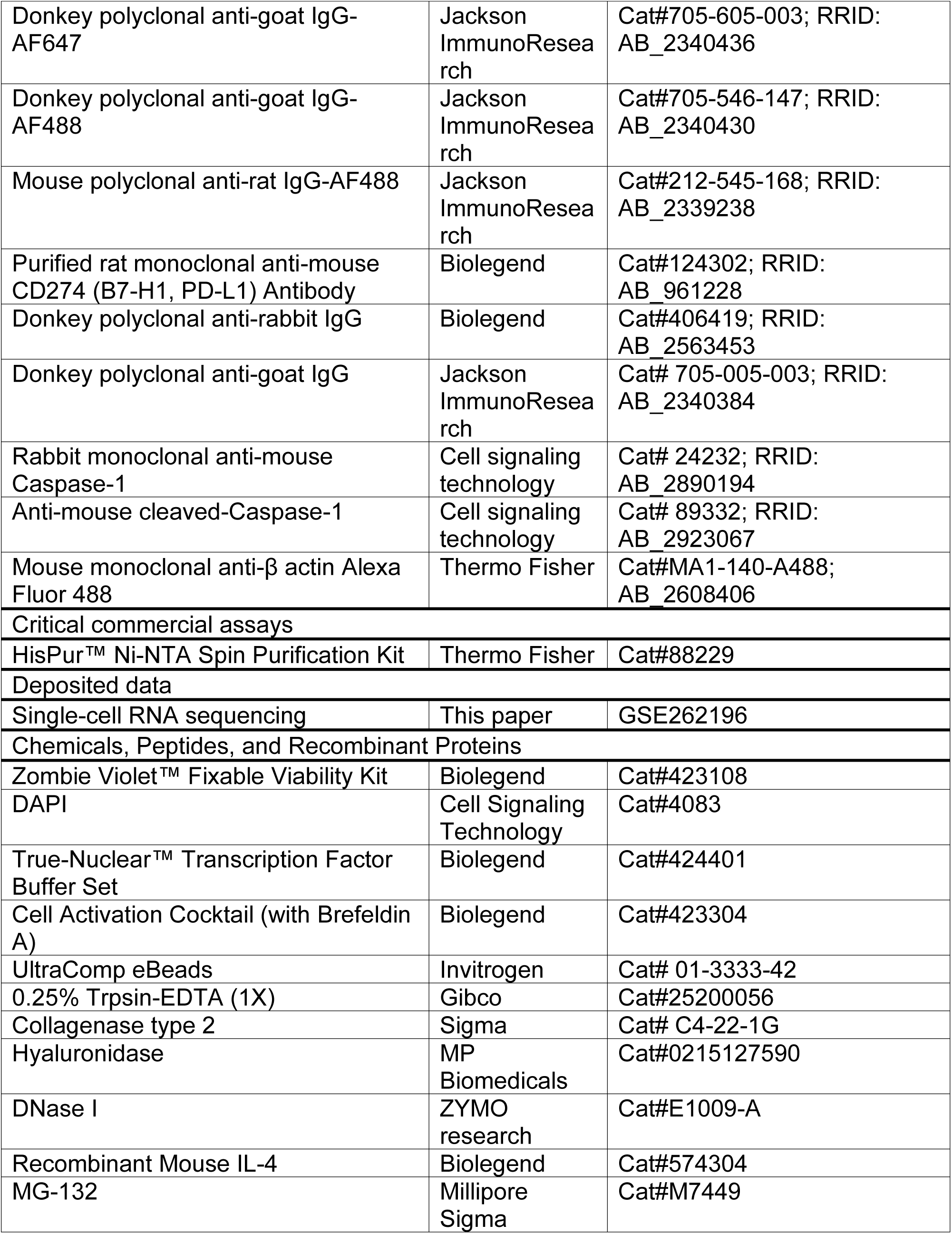
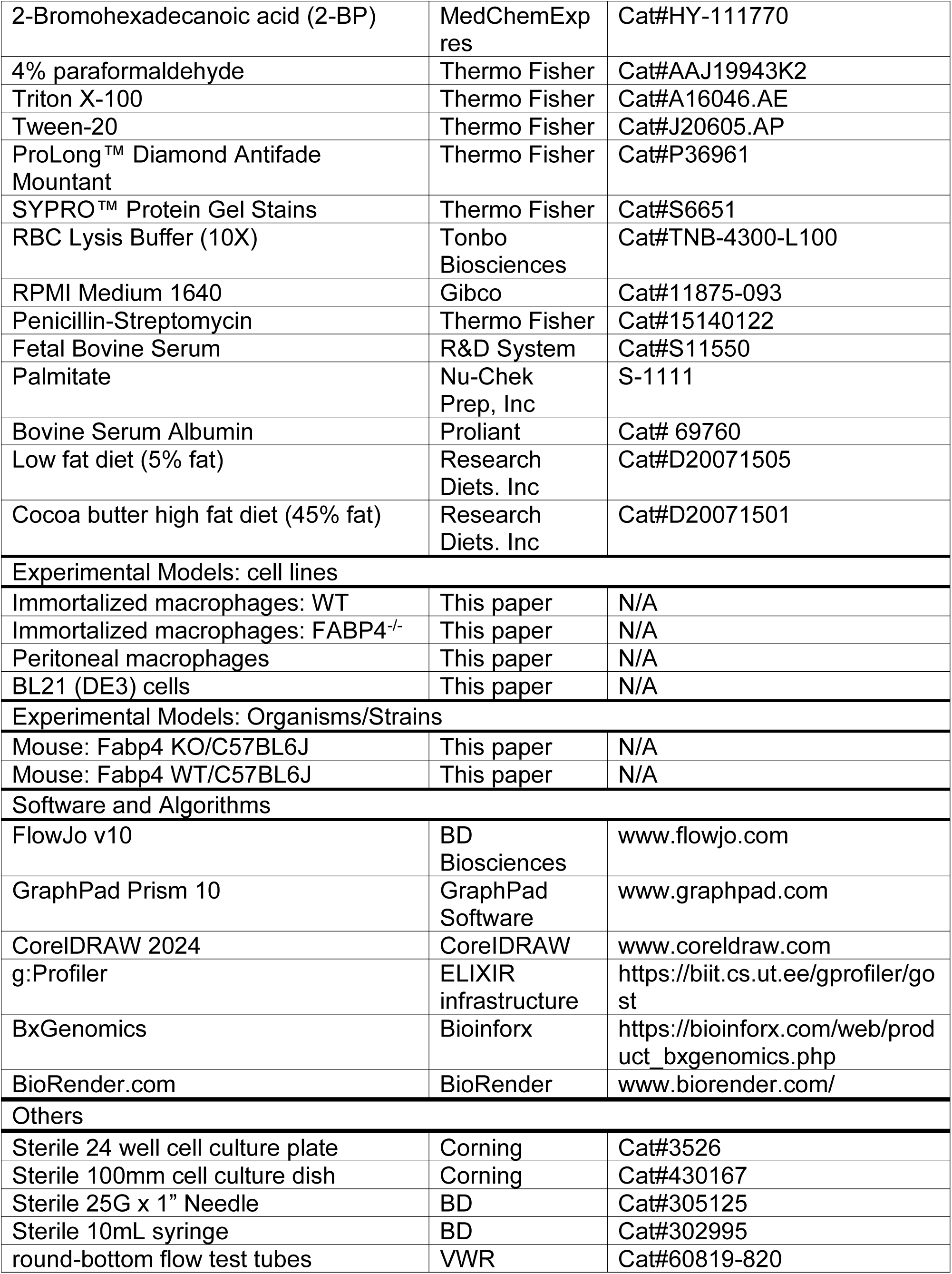
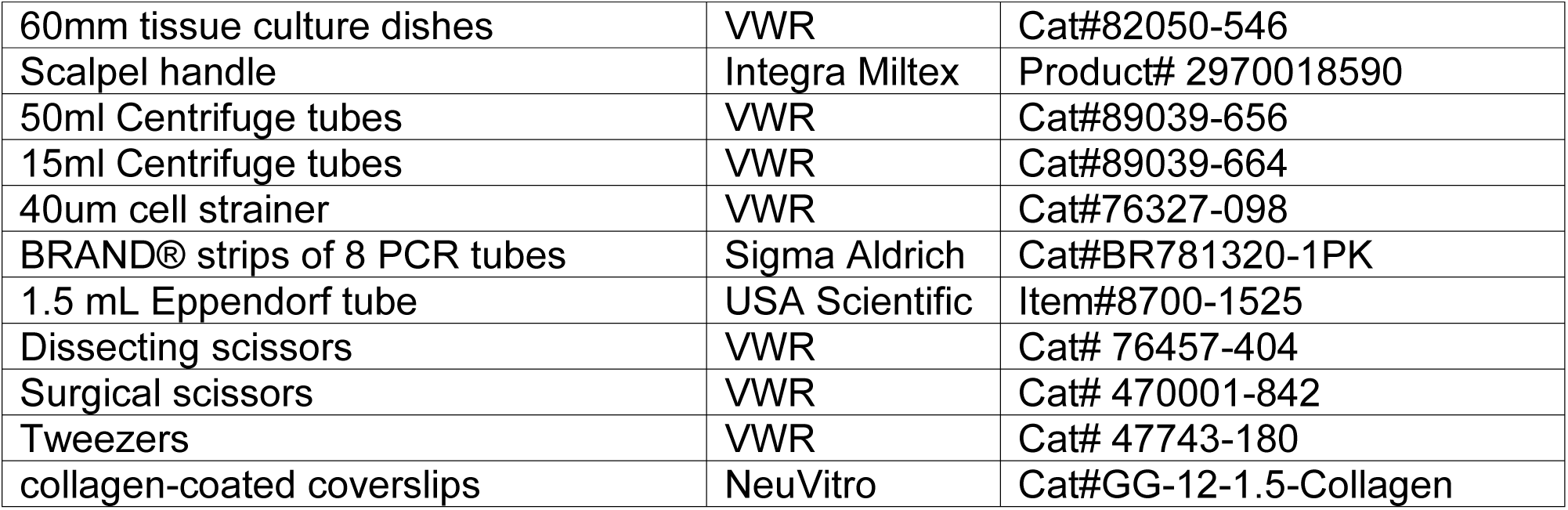

### EXPERIMENTAL MODEL AND STUDY PARTICIPANT DETAILS

#### Mice and syngeneic breast cancer model

The protocol of mouse studies was approved by the Institutional Animal Care and Use Committee (IACUC) at the University of Iowa. *Fabp4*^-/-^ mice and their wild-type controls were bred and housed in the animal facility at the University of Iowa. Male and female mice were weaned at 3–4 weeks of age and randomly assigned to different groups for special diet feeding, which included a low-fat diet (LFD, 5% fat from soybean) and a high-fat diet (HFD, 40% fat from cocoa butter) for 3 months. 5 × 10^5^ E0771 cells were orthotopically implanted into the fat pad of the fourth mammary gland of lean or obese mice on different diets, respectively. Tumor growth was monitored at 3-day intervals with calipers and tumor volume was calculated by the formula 0.5 × (large diameter) × (small diameter)^2^. Tumors and lungs were weighed at the end point when mice were sacrificed.

#### Human Subjects

Patient samples were collected at the Molecular Epidemiology Resources, University of Iowa with approved IRB protocols (201003791, 202107133). Peripheral blood samples were collected and transported to the study team for biomarker screening without further patient identifiers. PBMCs were separated for further analysis.

## MODEL DETAILS

### Single-cell suspension preparation

For human peripheral blood monocytes (PBMCs), 5mL blood was mixed with equal volume of cold PBS then was added to the surface of 7.5mL Ficoll (Cytiva, 1.078g/mL). PBMCs were obtained by centrifugation at 400g for 30 minutes at room temperature and collecting the interface cell layer. Cells were washed with cold PBS, pelleted, and resuspended in cold PBS.

For mouse PBMCs, 400uL blood from mice were added into 5mL RBC lysis buffer (Tonbo Biosciences) and incubated at room temperature for 15 min. Cells were washed with cold PBS, pelleted, and process 15 min of RBC lysis once more. Cells were then washed again, pelleted, and resuspended in cold PBS.

For mouse splenic single-cells, mouse spleen was collected and ground to release single cells. Cells were filtered through a 40μm cell stainer and pelleted. Cells were then processed red blood cell lysis by resuspending in 3mL RBC lysis buffer (Tonbo Biosciences) and incubated for 3 min at room temperature. Cells were washed with cold PBS after RBC lysis, pelleted, and resuspended in cold PBS.

For tumor single cells, EO771 tumor mass were collected, cut into small pieces, and digested by tri-enzyme digestion buffer (0.02mg/mL DNase I, 0.5mg/mL Collagenase type 2, 0.2 mg/mL Hyaluronidase) at 25rpm in 37°C for 45 minutes. Tumor single cells were obtained by vertexing tissue for 20 seconds and filtered through a 40μm cell stainer and pelleted. Cells were washed with cold PBS, pelleted, and resuspended in cold PBS.

#### Single-cell RNA sequencing

Splenic macrophages from mice fed different special diets were used for the 10 x Genomics 3’ expression single cell assay. Briefly, splenocytes were isolated from 3 pairs of age-matched WT female mice. Anti-CD16/32 antibody was used to block Fc receptors before additional staining with Zombie-violet (live/death dye), anti-mouse F4/80 antibody and anti-mouse CD11b antibody. The F4/80+/CD11b+/Zombie-Voilet-cells were sorted using a FACS Aira III instrument and resuspended at a concentration of 1000 cell/μL cold PBS with 0.04% non-acetylated BSA. An equal number of 5000 targeted cells from each sample were prepared to create GEMs. The GEM generation/barcoding, post GEM-reverse transcription cDNA amplification, and library construction were performed according to the manufacturer’s guidelines.

#### Single-cell data analysis

Single-cell gene counting was performed by Cell Ranger (10 X Genomics, version 7.1.0) using the refdata-gex-mm10-2020-A reference transcriptome. The resulting data matrices were subsequently imported into R (version 4.2.3) and analyzed using the Seurat package (version 4.9.9). Cells with less than 200 features or with percent mitochondrial gene expression greater than 5% were excluded from the analysis. The data across samples was integrated using the Integrate Data function. Gene expression was normalized and scaled using the default parameters. Based on visual inspection of the elbow plot, the first 30 Principal Components (PC) were used in UMAP-based dimensional reduction. The FindClusters function, with a resolution of 0.5, was then used to assign cells to clusters. The FindMarkers function was used to identify genes differentially expressed between HFD and LFD groups or between different clusters within the same sample type. Processed single-cell data sets were visualized using the BxGenomics scRNA-Seq View (https://app.bxgenomics.com/bxg/app/).

#### Flow cytometry

For mouse single cell surface staining, cells were incubated in cold PBS for 30 min with the following flow antibodies: Zombie-violet(for live/death cells), anti-mouse CD45-BUV563, CD11b-BUV737, TCR γ/δ-PE-Cy5, CD4-PE-Fire700, CD8α-RB613, B220-AF700, F4/80-AF488, CD11c-BV711, NK1.1-BV650, MHCII-BV605, Ly6C-BB700, Ly6G-Spark blue-550, CD3-RB744, PD-L1-PE-Cy7, and CD16/32 (Fc blocker). For intracellular FABP4 staining, cells stained by surface markers underwent the intracellular staining process using anti-mouse FABP4 antibody for 1.5 hours at 4°C, followed by secondary staining of Donkey-anti-goat IgG-AF647 antibody. Cells were acquired by Cytek Aurora flow cytometer.

For human PBMC surface staining, cells were incubated in cold PBS for 30 min with the following flow antibodies: Zombie-violet (for live/death cells), anti-human CD3-BUV737, CD4-AF700, CD8-PE-Dazzle594, CD19-Percp-Cy5.5, TCR γ/δ-PE-Fire700, CD14-Pacific Blue, CD16-BV650, CD56-BUV805, PD-L1-BV785. For intracellular cytokine staining, cells were treated with a Cell Activation Cocktail (Biolegend in RPMI-1640 medium containing 5% fetal bovine serum (FBS) for 4 h at 37°C, followed by the surface staining using the above anti-human antibodies. Afterward, cells were washed with PBS, fixed, and permeabilized using the intracellular staining buffer set (Cat. #424401, Biolegend) according to the manufacturer’s instructions. Intracellular staining was performed using the following antibodies: anti-human TNFα-PE-Cy7, IL-2-APC-Cy7, IFNγ-AF647. For intracellular FABP4 staining, cells stained by surface markers underwent the intracellular staining process using anti-human FABP4 antibody for 1.5 hours at 4°C, followed by secondary staining of Donkey-anti-goat IgG-AF488 antibody. Cells were acquired by Cytek Aurora flow cytometer.

#### Cell culture

Immortalized macrophage cell lines were established from Fabp4^-/-^ (FABP4^-/-^macrophages) or WT mice (WT macrophages) as described previously^51, 52^. For M2 polarization and PD-L1 detection, WT and FABP4^-/-^ macrophage were treated with recombinant mouse IL-4 (20ng/mL) in RPMI-1640 containing 10% FBS for 16 hours. Medium was discarded and replaced with fresh medium without IL-4. MG132 (2uM) or DMSO control added and continue culture for another 6 hours. Surface PD-L1 were detected by flow cytometry at timepoints of 0 hours, 16 hours, 19 hours, and 22 hours.

#### Fatty acid (FFA) preparation and treatment

Palmitic acids (PA) were conjugated with bovine serum albumin (BSA) as described ^28, 53^. Briefly, PA was conjugated with a pre-prepared solution of 2 mM endotoxin-free, fatty acid-free BSA in PBS at a concentration of 5 mM. The PA-BSA conjugates were sonicated until fully dissolved, then filtered through a 0.2 μm sterile filter for use in cell culture studies. Conjugates were subsequently added to the cell culture system at the specified final concentration of 400μM in RPMI-1640 containing 10% FBS.

#### FABP4 protein purification and thermal shift assay

Recombinant His_6_-tagged FABP4 was expressed in BL21 (DE3) cells and purified using Ni-NTA affinity chromatography. The His₆ tag was subsequently removed by TEV protease digestion, followed by further purification via ion-exchange chromatography. To eliminate residual endogenous lipids, the protein underwent a final purification step using a Lipidex-1000 column, yielding lipid-free (apo) FABP4.

The purified apo FABP4 protein was then used to assess fatty acid interactions in a fluorescence based thermal shift assay ^54^. Reactions were prepared in PCR tubes with a total volume of 20 μl, containing 10 μM FABP4 and 10× SYPRO Orange dye (Invitrogen) in buffer (20 mM HEPES, pH 7.0, 150 mM NaCl), along with either test compounds or ethanol controls. Samples were sealed, briefly centrifuged, and subjected to a temperature gradient from 25°C to 95°C at a rate of 1°C per minute using a 7500 Real-Time PCR system (Applied Biosystems). Raw fluorescence data were analyzed and melting temperatures (Tm) were determined through curve fitting as previously described.

#### H&E staining

Fresh lung tissues were obtained from euthanized mice. Lung samples were fixed immediately in 10% neutral buffered formalin for 24 h. Following serial alcohol dehydration (50%, 75%, 95%, and 100%), the samples were embedded in paraffin. The paraffin-embedded samples were sliced into 5 μm sections and stained in the DRS-601 Auto Stainer with hematoxylin and eosin (H&E) for 1 min. Slides were mounted with VectaMount Express Mounting Medium (vector laboratories, H-5700-60) and were scanned by slide scanner (Leica Aperio GT 450) for quantification analysis. The metastatic tumor number was analyzed by the SlideViewer 2.7.0.191696 software.

#### Confocal microscopy

Immortalized macrophage cell lines were seeded onto collagen-coated coverslips (NeuVitro; #GG-12-1.5-Collagen) at a density of 5 × 105 cells per well in a 24-well plate. Two hours after cell adhesion, 400 μM palmitic acid (PA) was added for the indicated time points. For surface PD-L1 staining, cells were washed twice with 1× PBS and incubated with rat anti-mouse PD-L1 antibody (1:50; BioLegend; #124301) diluted in RPMI at 37 °C for 7 min. Cells were then fixed with 4% paraformaldehyde for 25 min and permeabilized with 0.5% Triton X-100 for 15 min. For intracellular PD-L1 staining, cells were fixed and permeabilized immediately after fatty acid treatment without prior surface staining. To block nonspecific binding, cells were incubated in blocking solution containing 1% BSA and 0.1% Tween-20 for 1 h. Primary antibodies, including rat anti-PD-L1 and goat anti-FABP4 (1:300; R&D Systems; #AF1443), were diluted in blocking solution and incubated overnight at 4 °C. Secondary antibodies, including mouse anti-rat Alexa Fluor® 488 (1:1000; Jackson ImmunoResearch; #212-545-168), donkey anti-goat Alexa Fluor® 647 (1:1000; Jackson ImmunoResearch; #705-605-003), and DAPI (1:2000; Cell Signaling Technology; #4083), were incubated in 1% BSA for 2 h at room temperature. Samples were washed three times with 1× PBS for 5 min between each step. Coverslips were mounted using ProLong Diamond Antifade Mountant (ThermoFisher; #P36961). Images were acquired using a Zeiss LSM 880 confocal microscope.

#### Randomization and Blinding

*In vivo* experiments: Adult mice (8–10 weeks old) were randomly assigned to dietary and treatment groups. Investigators were blinded to group identity during tumor measuring and histological assessment. *In vitro* experiments: Treatment conditions were randomly assigned to wells to avoid positional bias, and replicate cultures were processed in parallel using blinded treatment labels.

## Quantification and statistical analysis

All data were presented as the mean ± SD. All experiments were performed by at least three independent replicates. For both *in vitro* and *in vivo* experiments, two-tailed, unpaired or paired Student’s t-test, two-tailed unpaired multiple t test, and one-way ANOVA with Bonferroni’s multiple comparison, two-way ANOVA with Tukey’s multiple comparison were used. *, *P* < 0.05; **, *P* < 0.01; ***, *P* < 0.001; ****, *P* < 0.0001; ns, nonsignificant. Linear regression was performed to assess the relationship between variables, and goodness-of-fit was evaluated using the coefficient of determination (R²) along with 95% confidence intervals (CI). Statistical significance was defined as *p* values.

