## Supplemental materials for "FABP4 Couples Lipid Metabolism to PD-L1 Stabilization in Immunosuppressive Macrophages"

**Supplemental Figure S1-S9**

**Supplemental Table S1**

### Figure-S1

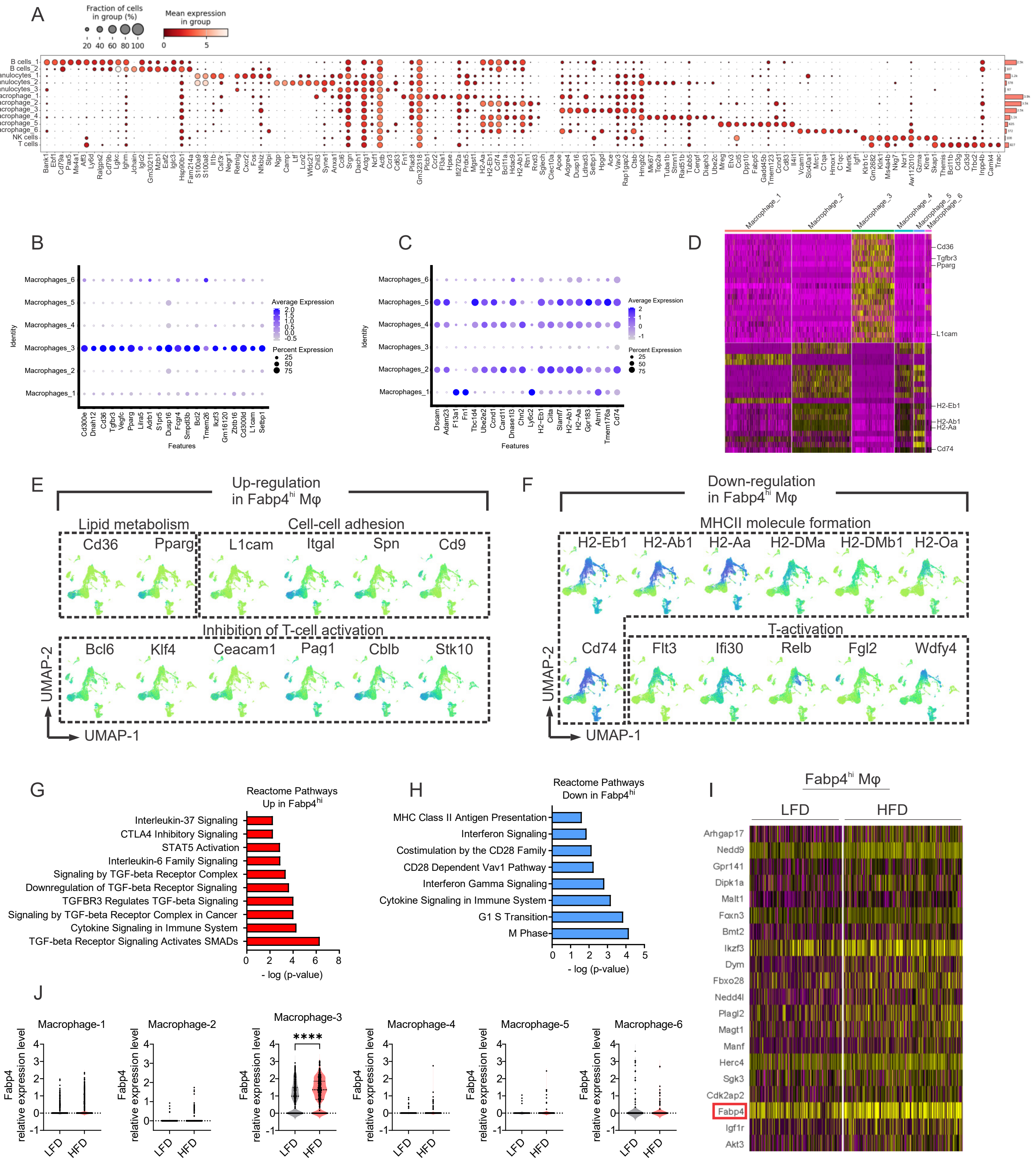
A
Fraction of cells in group (%)
20 40 60 80 100
Mean expression in group
0 5
B cells_1
B cells_2
Granulocytes_1
Granulocytes_2
Granulocytes_3
Macrophage_1
Macrophage_2
Macrophage_3
Macrophage_4
Macrophage_5
Macrophage_6
NK cells
T cells
B
Macrophages_6
Macrophages_5
Macrophages_4
Macrophages_3
Macrophages_2
Macrophages_1
Identity
Average Expression
Percent Expression
25
50
75
Cd300e
Dnah12
Cd36
Tgfbr3
Vegfc
Pparg
Lilra5
Adrb1
S1pr5
Dusp16
Fcgr4
Smpdl3b
Bcl2
Tmem26
Ikzf3
Gm16120
Zbtb16
Cd300ld
L1cam
Selbp1
Features
C
Dscam
Adam23
F13a1
Fn1
Tbc1d4
Ube2e2
Ccnd1
Card11
Dnase1l3
Ctnn2
Ly6c2
H2-Eb1
Ciita
Slamf7
H2-Ab1
H2-Aa
Gpr183
Atrnl1
Tmem176a
Cd74
Features
D
Macrophage_1
Macrophage_2
Macrophage_3
Macrophage_4
Macrophage_5
Macrophage_6
Cd36
Tgfbr3
Pparg
L1cam
H2-Eb1
H2-Ab1
H2-Aa
Cd74
E
Up-regulation in Fabp4hi Mφ
Lipid metabolism
Cell-cell adhesion
Cd36
Pparg
L1cam
Itgal
Spn
Cd9
Inhibition of T-cell activation
Bcl6
Klf4
Ceacam1
Pag1
Cblb
Stk10
UMAP-2
UMAP-1
F
Down-regulation in Fabp4hi Mφ
MHCII molecule formation
H2-Eb1
H2-Ab1
H2-Aa
H2-DMa
H2-DMb1
H2-Oa
T-activation
Cd74
Flt3
Ifi30
Relb
Fgl2
Wdfy4
UMAP-2
UMAP-1
G
Reactome Pathways Up in Fabp4hi
Interleukin-37 Signaling
CTLA4 Inhibitory Signaling
STAT5 Activation
Interleukin-6 Family Signaling
Signaling by TGF-beta Receptor Complex
Downregulation of TGF-beta Receptor Signaling
TGFBR3 Regulates TGF-beta Signaling
Signaling by TGF-beta Receptor Complex in Cancer
Cytokine Signaling in Immune System
TGF-beta Receptor Signaling Activates SMADs
0 2 4 6 8
- log (p-value)
H
Reactome Pathways Down in Fabp4hi
MHC Class II Antigen Presentation
Interferon Signaling
Costimulation by the CD28 Family
CD28 Dependent Vav1 Pathway
Interferon Gamma Signaling
Cytokine Signaling in Immune System
G1 S Transition
M Phase
0 1 2 3 4 5
- log (p-value)
I
Fabp4hi Mφ
LFD
HFD
Arhgap17
Nedd9
Gpr141
Dipk1a
Malt1
Foxn3
Bmt2
Ikzf3
Dym
Fbxo28
Nedd4l
Plagl2
Magt1
Manf
Herc4
Sgk3
Cdk2ap2
Fabp4
Igf1r
Akt3
J
Macrophage-1
Macrophage-2
Macrophage-3
Macrophage-4
Macrophage-5
Macrophage-6
Fabp4 relative expression level
****
LFD
HFD

**Figure S1 Single-cell RNA sequencing analysis of mouse F4/80$^{+}$-enriched splenocytes in LFD and HFD,** related to Figure 1

(A) Dot plot of canonical cell lineage-defining markers for the populations annotated in Figure 1A.

(B) Dot plot of top 20 up-regulated genes in macrophage_3 cluster compared to other macrophage clusters.

(C) Dot plot of top 20 down-regulated genes in macropohage_3 cluster compared to other macrophage clusters.

(D) Heatmap of top 20 DEGs in macrophage_3 cluster compared to other clusters.

(E) UMAPs of representative up-regulated genes in macrophage_3 cluster compared to other macrophage clusters.

(F) UMAPs of representative down-regulated genes in macrophage_3 cluster compared to other macrophage clusters.

(G) Pathway enrichment analysis of significantly upregulated genes in Fabp4$^{hi}$ compared to Fabp4$^{lo}$ splenic macrophages using Reactome Pathways database.

(H) Pathway enrichment analysis of significantly downregulated genes in Fabp4$^{hi}$ compared to Fabp4$^{lo}$ splenic macrophages using Reactome Pathways database.

(I) Heatmap of top 20 up-regulated DEGs in macrophage_3 cluster from mice fed HFD compared LFD.

(J) Violin plots showing relative expression levels of Fabp4 across macrophage subsets in scRNA-seq.

### Figure-S2

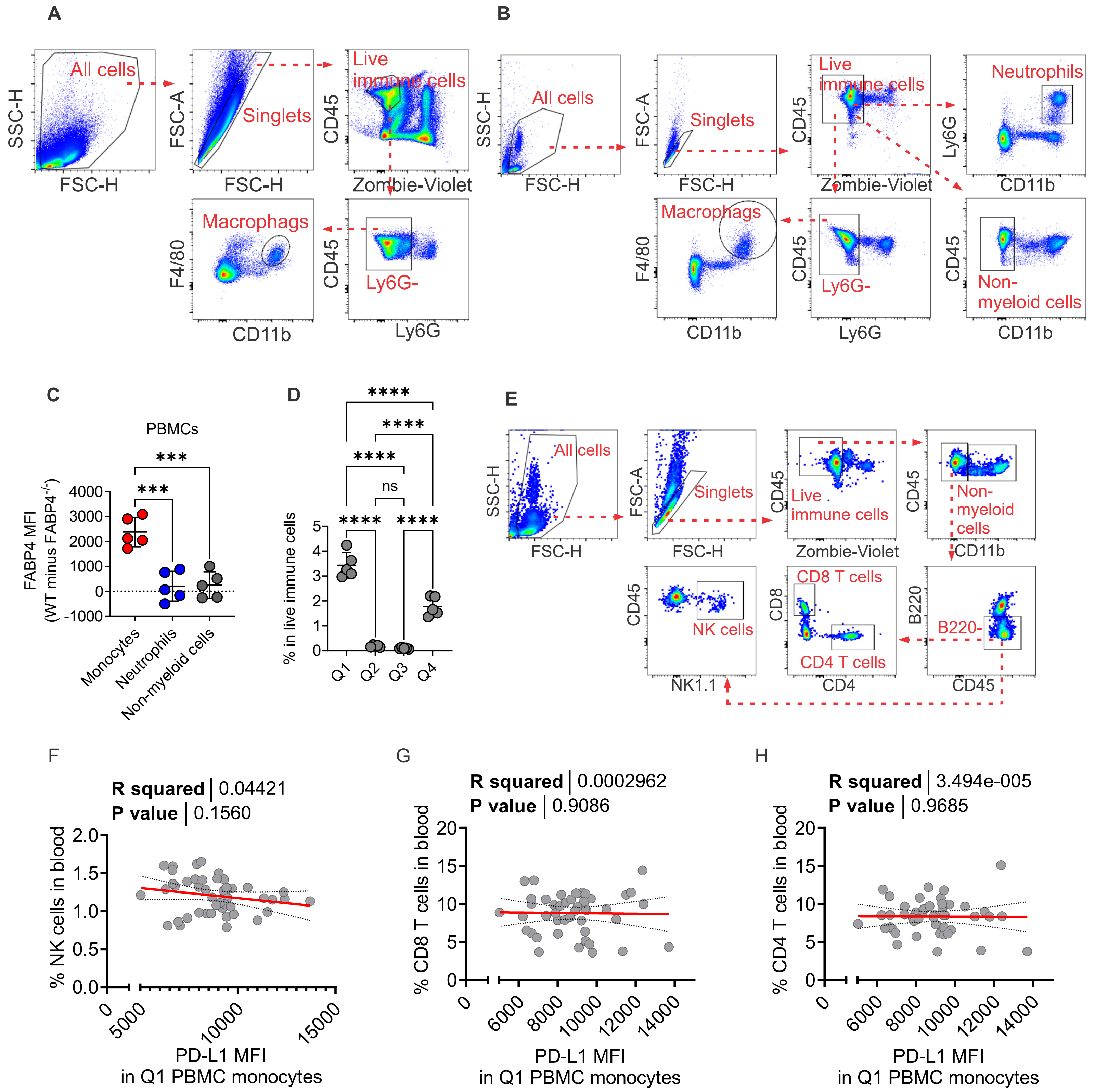

A
All cells
SSC-H
FSC-H
Singlets
FSC-A
Live immune cells
CD45
Zombie-Violet
Macrophags
F4/80
CD11b
Ly6G-
Ly6G
B
Neutrophils
Ly6G
CD11b
Non-myeloid cells
C
PBMCs
FABP4 MFI (WT minus FABP4-/-)
***
Monocytes
Neutrophils
Non-myeloid cells
D
% in live immune cells
****
ns
Q1
Q2
Q3
Q4
E
NK cells
NK1.1
CD8 T cells
CD4 T cells
CD8
CD4
B220-
B220
F
R squared | 0.04421
P value | 0.1560
% NK cells in blood
PD-L1 MFI in Q1 PBMC monocytes
G
R squared | 0.0002962
P value | 0.9086
% CD8 T cells in blood
H
R squared | 3.494e-005
P value | 0.9685
% CD4 T cells in blood

**Figure S2 Analysis of PD-L1 expression in PBMC monocytes,** related to Figure 2.

(A) Proposed gating strategy for mouse splenic macrophage by flow cytometry.

(B) Proposed gating strategy for mouse PBMC monocytes, neutrophils, and non-myeloid cells by flow cytometry.

(C) Flow cytometric analysis of intracellular FABP4 levels across PBMC immune cell subsets, shown as representative flow plots and quantified as relative intracellular FABP4 MFI by subtracting the $FABP4^{-/-}$ from WT.

(D) Flow cytometric analysis of monocyte subset (Q1-Q4) ratios in live immune cells in PBMC.

(E) Proposed gating strategy for mouse PBMC NK cells, $CD8^{+}$ T cells, and $CD4^{+}$ T cells by flow cytometry.

(F) Correlation analysis of surface PD-L1 MFI in Q1 PBMC monocyte subset versus NK cell percentage in PBMC by flow cytometry

(G) Correlation analysis of surface PD-L1 MFI in Q1 PBMC monocyte subset versus $CD8^{+}$ T cell percentage in PBMC by flow cytometry.

(H) Correlation analysis of surface PD-L1 MFI in Q1 PBMC monocyte subset versus $CD4^{+}$ T cell percentage in PBMC by flow cytometry.

Data are presented as the mean ± SD. *, $P < 0.05$; **, $P < 0.01$; ****, $P < 0.0001$; ns, nonsignificant; one-way ANOVA with Bonferroni’s multiple comparison test for (C) and (D), linear regression analysis for (F-H).

### Figure-S3

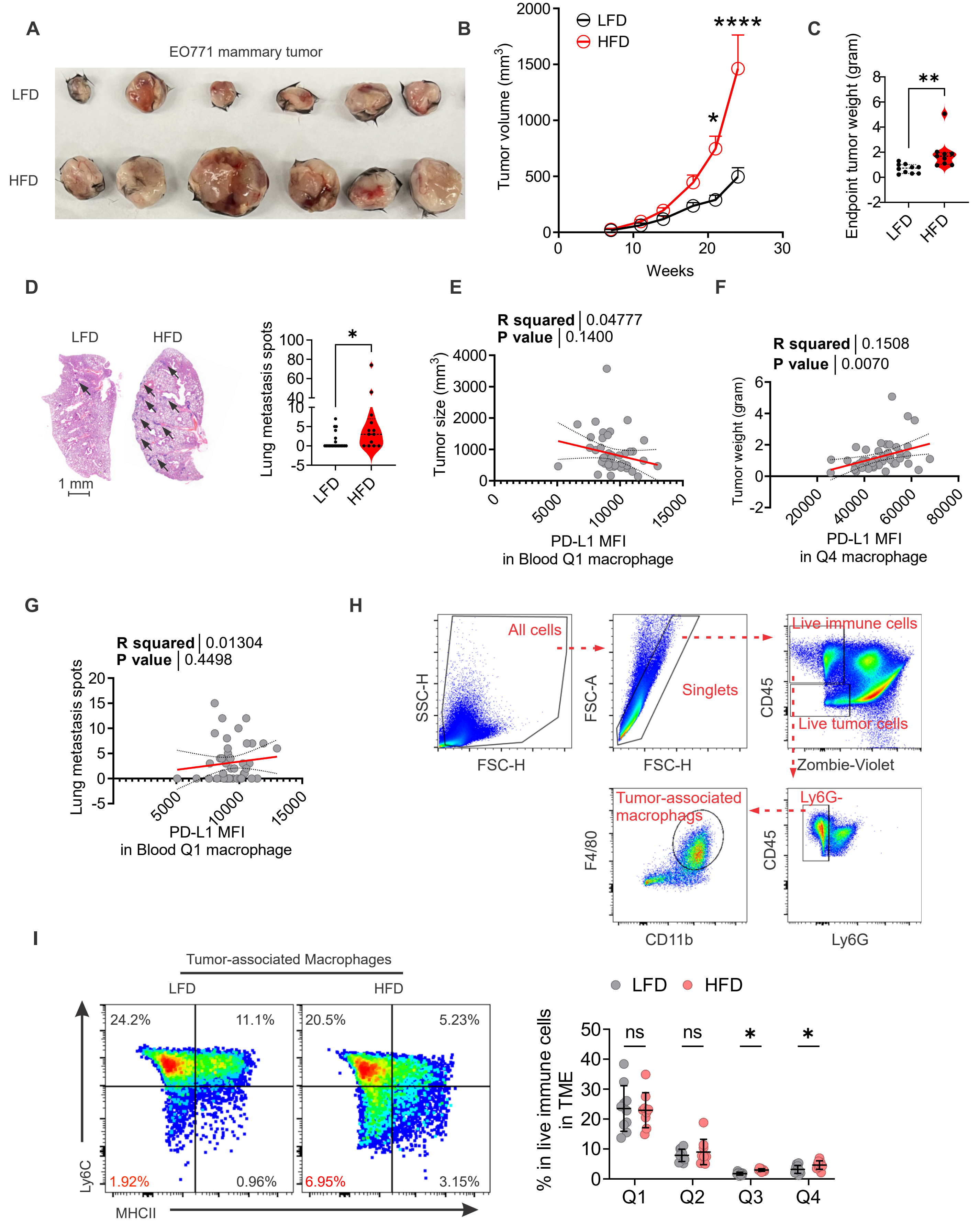

A
EO771 mammary tumor
LFD
HFD
B
LFD
HFD
****
*
Tumor volume (mm³)
Weeks
C
**
Endpoint tumor weight (gram)
LFD
HFD
D
LFD
HFD
1 mm
*
Lung metastasis spots
LFD
HFD
E
R squared | 0.04777
P value | 0.1400
Tumor size (mm³)
PD-L1 MFI
in Blood Q1 macrophage
F
R squared | 0.1508
P value | 0.0070
Tumor weight (gram)
PD-L1 MFI
in Q4 macrophage
G
R squared | 0.01304
P value | 0.4498
Lung metastasis spots
PD-L1 MFI
in Blood Q1 macrophage
H
All cells
SSC-H
FSC-H
Singlets
FSC-A
FSC-H
Live immune cells
Live tumor cells
CD45
Zombie-Violet
Tumor-associated macrophags
F4/80
CD11b
Ly6G-
CD45
Ly6G
I
Tumor-associated Macrophages
LFD
HFD
24.2%
11.1%
1.92%
0.96%
20.5%
5.23%
6.95%
3.15%
Ly6C
MHCII
LFD
HFD
% in live immune cells in TME
ns
ns
*
*
Q1
Q2
Q3
Q4

**Figure S3 HFD increased breast cancer burden in mice,** related to Figure 3**.**

(A) Representative Images showing endpoint tumor size from mice fed HFD and LFD, received orthotopically injection of EO771 tumor cells.

(B) Growth curve of tumor from mice fed HFD and LFD post orthotopically injection of EO771 tumor cells.

(C) Endpoint tumor weight from mice fed HFD and LFD post orthotopically injection of EO771 tumor cells.

(D) Metastasis burden of mice fed HFD and LFD, received orthotopically injection of EO771 tumor cells, shown as representative images of lung metastasis (black arrows) using histopathologic hematoxylin and eosin (H&E) staining, and quantified number of metastasis spots per lung slide.

(E) Correlation analysis of surface PD-L1 MFI by flow cytometry in Q1 PBMC monocyte subset versus endpoint tumor size after orthotopically injection of EO771 tumor cells into fourth mammary fat pad.

(F) Correlation analysis of surface PD-L1 MFI by flow cytometry in Q1 PBMC monocyte subset versus endpoint tumor weight after orthotopically injection of EO771 tumor cells into fourth mammary fat pad.

(G) Correlation analysis of surface PD-L1 MFI by flow cytometry in Q1 PBMC monocyte subset and number of lung metastasis spots after orthotopically injection of EO771 tumor cells into fourth mammary fat pad.

(H) Proposed gating strategy for tumor-associated macrophages in the EO771 tumor microenvironment (TME) by flow cytometry.

(I) Flow cytometric analysis of ratios of tumor-associated macrophage subsets in EO771 tumor tissue in LFD and HFD, shown as representative and the quantified ratios in live immune cells in the EO771 TME.

Data are presented as the mean ± SD. *, $P < 0.05$; **, $P < 0.01$; ****, $P < 0.0001$; ns, nonsignificant; two-way ANOVA with Tukey's multiple comparison test for (B), unpaired two-tailed t test for (C) and (D), unpaired two-tailed multiple t-test for (I), and linear regression analysis for (E), (F), and (G).

### Figure-S4

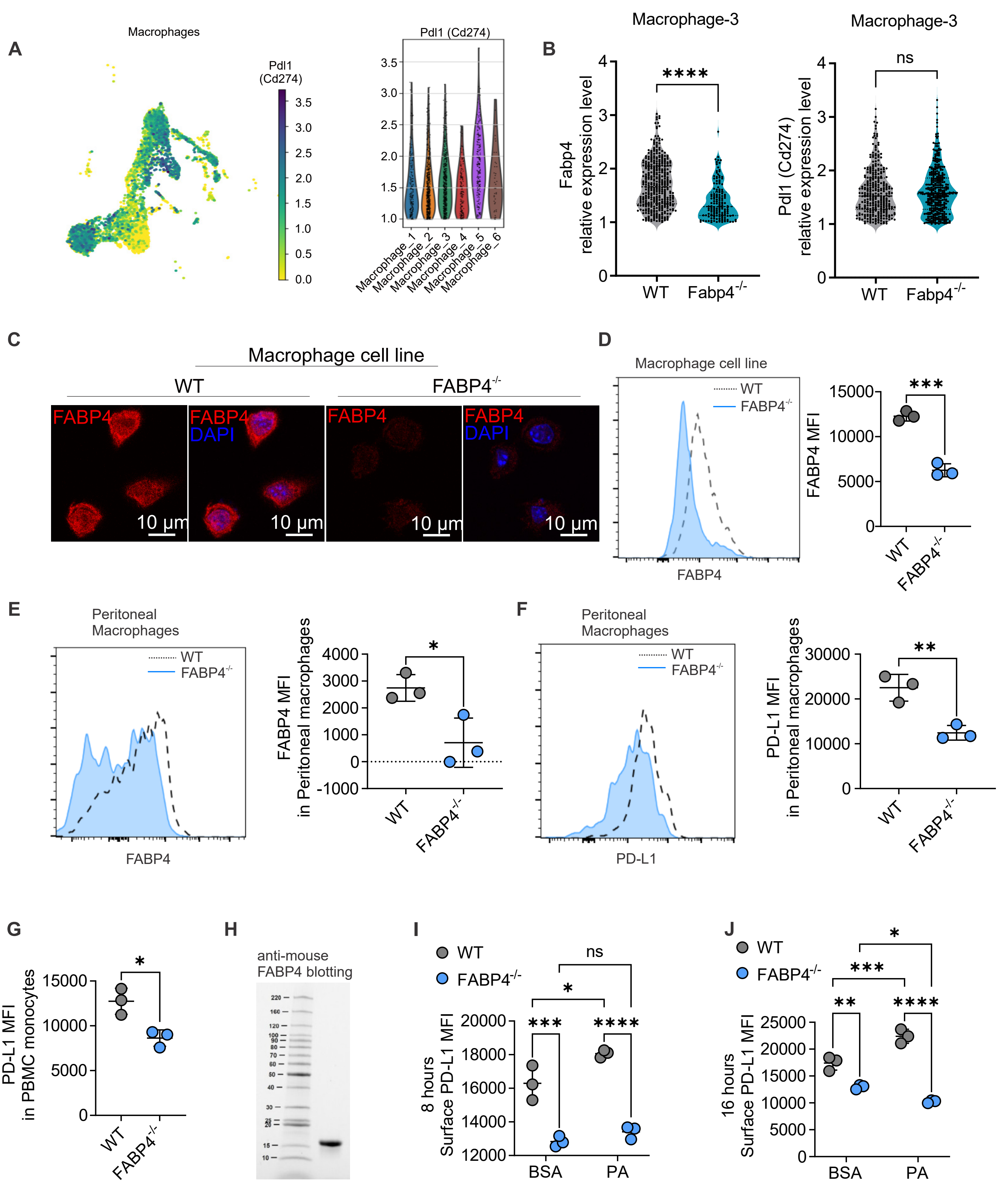

A
Macrophages
Pdl1 (Cd274)
Pdl1 (Cd274)
Macrophage_1
Macrophage_2
Macrophage_3
Macrophage_4
Macrophage_5
Macrophage_6
B
Macrophage-3
Fabp4 relative expression level
****
WT
Fabp4-/-
Macrophage-3
Pdl1 (Cd274) relative expression level
ns
WT
Fabp4-/-
C
Macrophage cell line
WT
FABP4-/-
FABP4
FABP4 DAPI
FABP4
FABP4 DAPI
10 μm
D
Macrophage cell line
WT
FABP4-/-
FABP4
FABP4 MFI
***
WT
FABP4-/-
E
Peritoneal Macrophages
WT
FABP4-/-
FABP4
FABP4 MFI in Peritoneal macrophages
*
WT
FABP4-/-
F
Peritoneal Macrophages
WT
FABP4-/-
PD-L1
PD-L1 MFI in Peritoneal macrophages
**
WT
FABP4-/-
G
PD-L1 MFI in PBMC monocytes
*
WT
FABP4-/-
H
anti-mouse FABP4 blotting
I
WT
FABP4-/-
8 hours Surface PD-L1 MFI
ns
*
***
****
BSA
PA
J
WT
FABP4-/-
16 hours Surface PD-L1 MFI
*
***
**
****
BSA
PA

**Figure S4 Intracellular FABP4 is required for macrophage PD-L1 surface expression,** related to Figure 4.

(A) Comparison of Pdl1 (Cd274) gene expression in across macrophage subsets in scRNA-seq, shown in UMAP and violin plots of relative expression.

(B) Violin plots of Fabp4 and Pdl1 (Cd274) gene relative expressions in macrophage-3 cluster in WT and Fabp4$^{-/-}$ mice in scRNA-seq.

(C) Validation of FABP4 deficiency in WT and FABP4$^{-/-}$ macrophage cell line using immunofluorescence staining.

(D) Validation of FABP4 deficiency in WT and FABP4$^{-/-}$ macrophage cell line using flow cytometry, shown in representative flow plots and quantified intracellular FABP4 MFI.

(E) Validation of FABP4 deficiency in peritoneal macrophages from WT and FABP4$^{-/-}$ mice using flow cytometry, shown in representative flow plots and quantified intracellular FABP4 MFI.

(F) Surface PD-L1 levels in WT and FABP4$^{-/-}$ peritoneal macrophage by flow cytometry, shown as representative flow plot and quantified surface PD-L1 MFI.

(G) Surface PD-L1 levels in PBMC monocytes from WT and FABP4$^{-/-}$ mice by flow cytometry, shown as representative flow plot and quantified surface PD-L1 MFI.

(H) Western blotting for purified recombinant mouse FABP4 protein.

(I) Quantified surface PD-L1 MFI in WT and FABP4$^{-/-}$ macrophages treated with BSA or PA for 8 hours in vitro by flow cytometry.

(J) Quantified surface PD-L1 MFI in WT and FABP4$^{-/-}$ macrophages treated with BSA or PA for 16 hours in vitro by flow cytometry.

Data are presented as the mean ± SD. *, $P < 0.05$; **, $P < 0.01$; ***, $P < 0.001$; ****, $P < 0.0001$; ns, nonsignificant; unpaired two-tailed t test for (B), (D), (E), (F), and (G), two-way ANOVA with Tukey's multiple comparison test for (I) and (J).

### Figure-S5

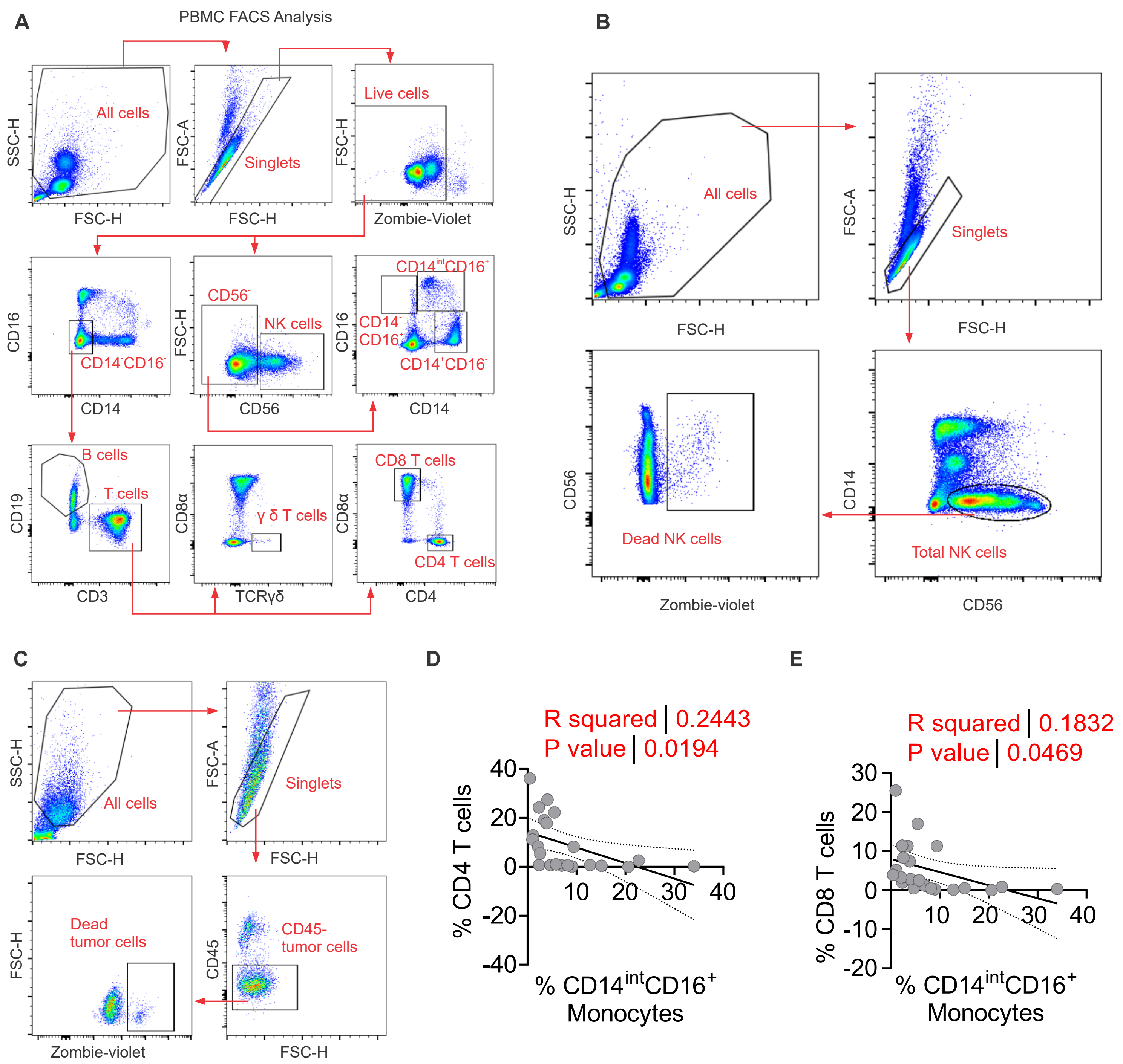

A
PBMC FACS Analysis
All cells
SSC-H
FSC-H
Singlets
FSC-A
FSC-H
Live cells
SSC-H
Zombie-Violet
CD16
CD14
CD14-CD16-
CD56-
NK cells
FSC-H
CD56
CD14intCD16+
CD14-CD16-
CD14+CD16-
CD16
CD14
B cells
T cells
CD19
CD3
γ δ T cells
CD8α
TCRγδ
CD8 T cells
CD4 T cells
CD8α
CD4
B
All cells
SSC-H
FSC-H
Singlets
FSC-A
FSC-H
Dead NK cells
CD56
Zombie-violet
Total NK cells
CD14
CD56
C
All cells
SSC-H
FSC-H
Singlets
FSC-A
FSC-H
Dead tumor cells
FSC-H
Zombie-violet
CD45- tumor cells
CD45
FSC-H
D
R squared | 0.2443
P value | 0.0194
% CD4 T cells
% CD14intCD16+ Monocytes
E
R squared | 0.1832
P value | 0.0469
% CD8 T cells
% CD14intCD16+ Monocytes

**Figure S5 Flow cytometric profiling of PBMC from non-cancer donors,** related to Figure 5.

(A) Proposed gating strategy for human PBMC subsets by flow cytometry.

(B) Proposed gating strategy for percentage of NK cell death in human PBMC by flow cytometry.

(C) Proposed gating strategy for percentage of MDA tumor cell death in coculture by flow cytometry.

(D) Correlation analysis of $CD14^{int}CD16^{+}$ monocyte ratio versus $CD4^{+}$ T cell ratio in coculture.

(E) Correlation analysis of $CD14^{int}CD16^{+}$ monocyte ratio versus $CD8^{+}$ T cell ratio in coculture.

Data are presented as the mean ± SD, linear regression analysis for (D) and (E).

### Figure-S6

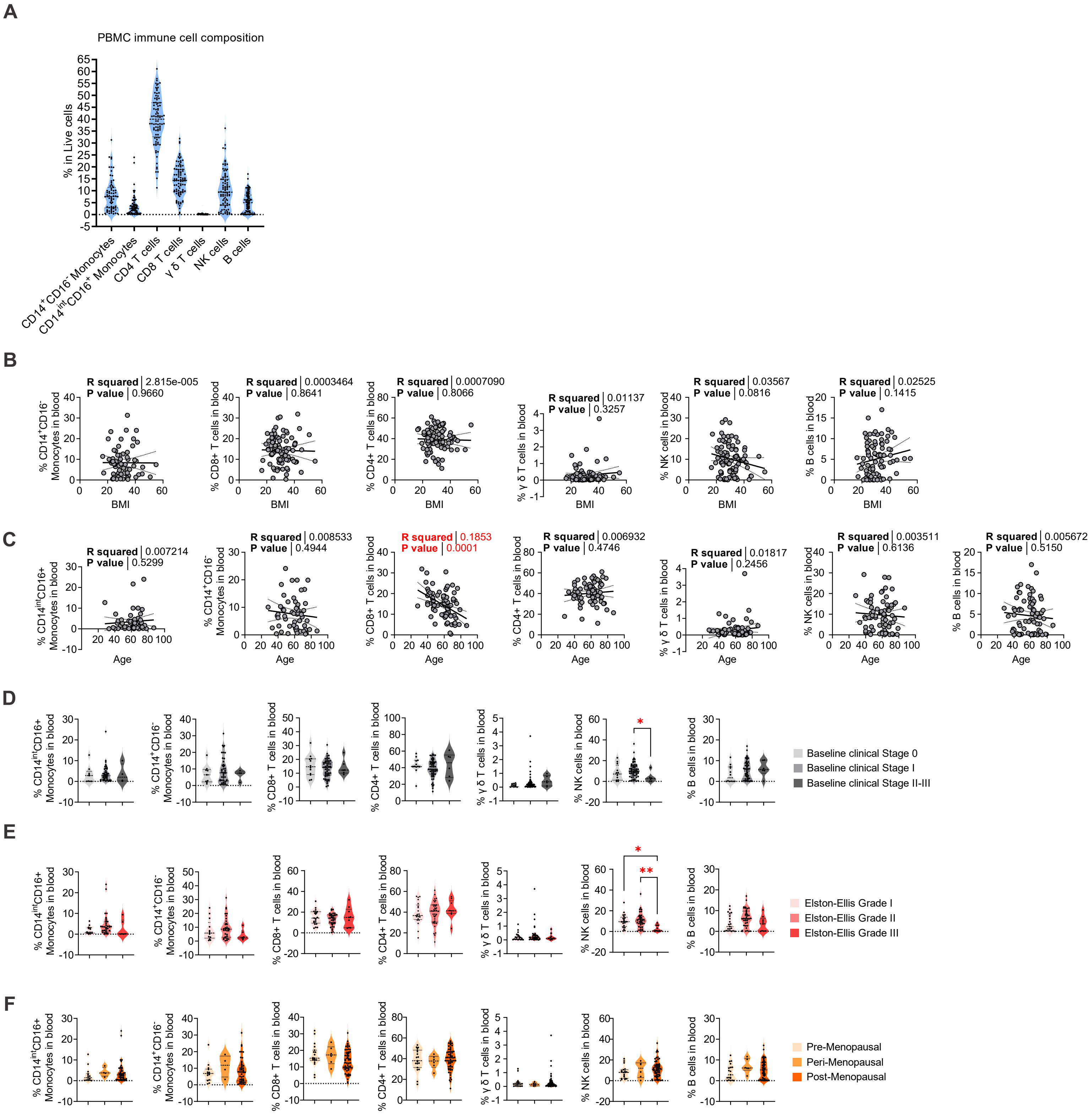
A
PBMC immune cell composition
% in Live cells
CD14+CD16- Monocytes
CD14intCD16+ Monocytes
CD4 T cells
CD8 T cells
γ δ T cells
NK cells
B cells
B
R squared 2.815e-005
P value 0.9660
R squared 0.0003464
P value 0.8641
R squared 0.0007090
P value 0.8066
R squared 0.01137
P value 0.3257
R squared 0.03567
P value 0.0816
R squared 0.02525
P value 0.1415
BMI
C
R squared 0.007214
P value 0.5299
R squared 0.008533
P value 0.4944
R squared 0.1853
P value 0.0001
R squared 0.006932
P value 0.4746
R squared 0.01817
P value 0.2456
R squared 0.003511
P value 0.6136
R squared 0.005672
P value 0.5150
Age
D
Baseline clinical Stage 0
Baseline clinical Stage I
Baseline clinical Stage II-III
E
Elston-Ellis Grade I
Elston-Ellis Grade II
Elston-Ellis Grade III
F
Pre-Menopausal
Peri-Menopausal
Post-Menopausal
% CD14intCD16+ Monocytes in blood
% CD14+CD16- Monocytes in blood
% CD8+ T cells in blood
% CD4+ T cells in blood
% γ δ T cells in blood
% NK cells in blood
% B cells in blood

**Figure S6 Correlations of PBMC subsets and clinical parameters in breast cancer patients,** related Figure 6**.**

(A) Breast cancer patient PBMC Immune profile shown as ratio of different subsets by flow cytometry.

(B) Correlation analysis of BMI versus ratios of different PBMC subsets in breast cancer patients.

(C) Correlation analysis of age versus ratios of different PBMC subsets in breast cancer patients.

(D) Ratios of different PBMC subsets in breast cancer patients across baseline clinical stages.

(E) Ratios of different PBMC subsets in breast cancer patients across histological Elston–Ellis's grades.

(F) Ratios of different PBMC subsets in breast cancer patients in different menopausal status.

Data are presented as the mean ± SD. *, $P < 0.05$; **, $P < 0.01$; ****, $P < 0.0001$; ns, nonsignificant; linear regression analysis for (B) and (C). one-way ANOVA with Bonferroni's multiple comparison test for (D-F).

### Figure-S7

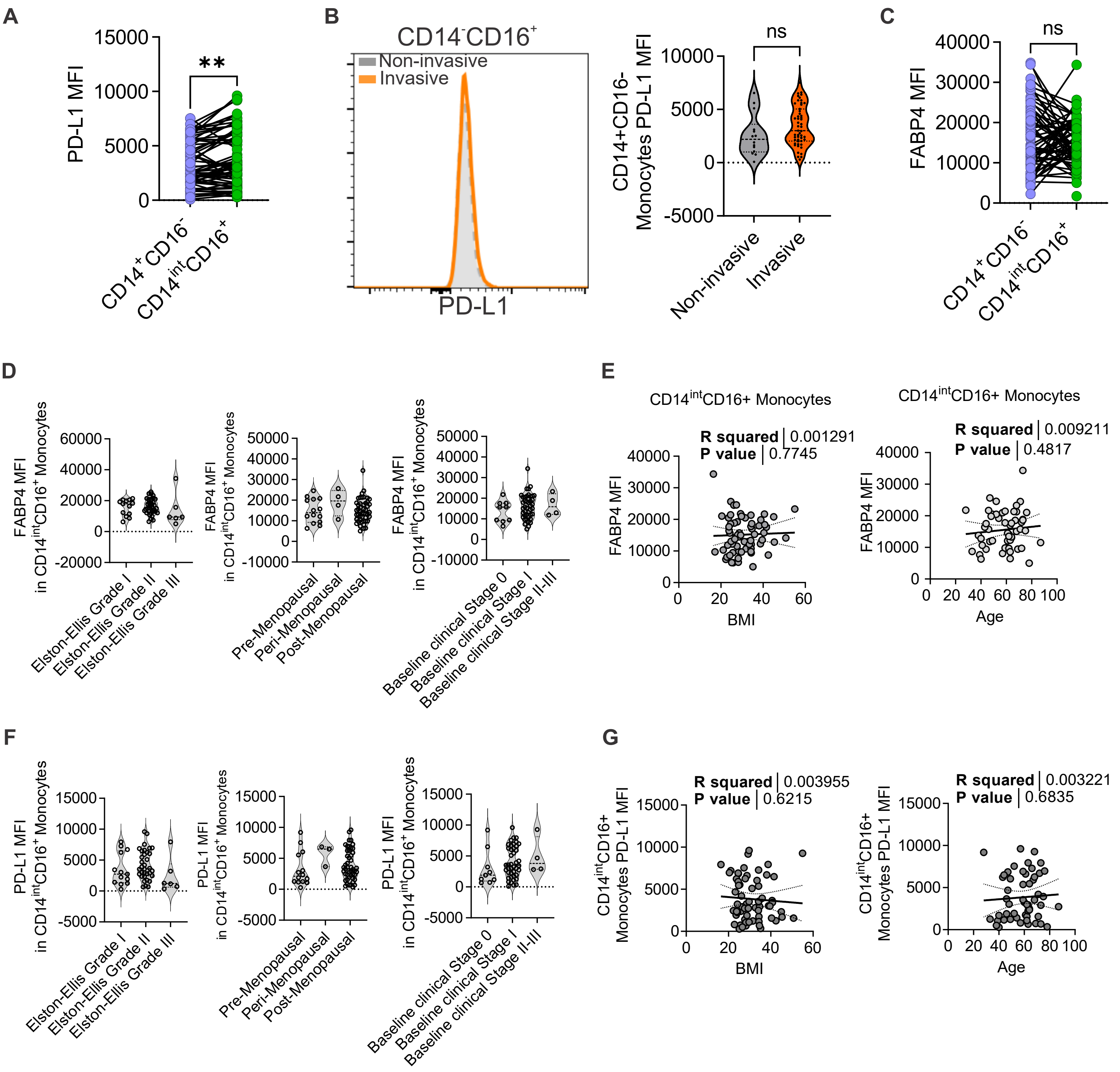

A
PD-L1 MFI
**
CD14+CD16-
CD14intCD16+
B
CD14-CD16+
Non-invasive
Invasive
PD-L1
CD14+CD16-
Monocytes PD-L1 MFI
ns
Non-invasive
Invasive
C
FABP4 MFI
ns
D
FABP4 MFI
in CD14intCD16+ Monocytes
Elston-Ellis Grade I
Elston-Ellis Grade II
Elston-Ellis Grade III
Pre-Menopausal
Peri-Menopausal
Post-Menopausal
Baseline clinical Stage 0
Baseline clinical Stage I
Baseline clinical Stage II-III
E
CD14intCD16+ Monocytes
R squared | 0.001291
P value | 0.7745
BMI
R squared | 0.009211
P value | 0.4817
Age
F
PD-L1 MFI
in CD14intCD16+ Monocytes
G
CD14intCD16+
Monocytes PD-L1 MFI
R squared | 0.003955
P value | 0.6215
R squared | 0.003221
P value | 0.6835

**Figure S7 Correlations of PD-L1 and FABP4 levels and clinical parameters in breast cancer patients,** related to Figure 7**.**

(A) Quantified surface PD-L1 MFI in $CD14^{+}CD16^{-}$ and $CD14^{int}CD16^{+}$ monocytes in breast cancer patients by flow cytometry.

(B) Surface PD-L1 levels in $CD14^{+}CD16^{-}$ monocytes in patients with non-invasive and invasive breast cancer types by flow cytometry, shown as representative flow plot and quantified surface PD-L1 MFI.

(C) Quantified intracellular FABP4 MFI in $CD14^{+}CD16^{-}$ and $CD14^{int}CD16^{+}$ monocytes by flow cytometry in breast cancer patients.

(D) Intracellular FABP4 MFI in $CD14^{int}CD16^{+}$ monocytes in breast cancer patients across histological Elston–Ellis's grades, menopausal status, and baseline clinical stages.

(E) Correlation analysis of BMI and age versus intracellular FABP4 MFI in $CD14^{int}CD16^{+}$ monocytes in breast cancer patients by flow cytometry.

(F) Quantified surface PD-L1 MFI in $CD14^{int}CD16^{+}$ monocytes by flow cytometry in breast cancer patients across histological Elston–Ellis's grades, menopausal status, and baseline clinical stages.

(G) Correlation analysis of BMI and age versus surface PD-L1 MFI in $CD14^{int}CD16^{+}$ monocytes in breast cancer patients by flow cytometry.

Data are presented as the mean ± SD. *, $P < 0.05$; **, $P < 0.01$; ****, $P < 0.0001$; ns, nonsignificant; paired two-tailed t-test for (A) and (C), unpaired two-tailed t-test for (B), one-way ANOVA with Bonferroni's multiple comparison test for (D) and (F). linear regression analysis for (E) and (G).

**Table S1. Clinical Patient Characteristics**

| | | |
|---|---|---|
| No. of Women Patient | 88 | |
| Age (yrs) | 58.5 ± 12.3 | |
| BMI (kg/m$^2$) | 30.6 ± 7.3 | |
| Waist Circumference (cm) | 100.0 ± 15.2 | |
| Histology | | |
| Invasive | 69 | 78% |
| IDC | 20 | 23% |
| ILC | 3 | 3% |
| Mixed | 46 | 52% |
| Non-invasive | 19 | 22% |
| DCIS | 10 | 11% |
| Other/mixed | 9 | 10% |
| Elston-Ellis Grade | | |
| 1 | 21 | 24% |
| 2 | 39 | 44% |
| 3 | 9 | 10% |
| Not Applicable | 19 | 22% |
| Clinical Stage | | |
| 0 | 12 | 14% |
| I | 58 | 66% |
| II | 5 | 6% |
| III | 1 | 1% |
| Not Applicable | 12 | 14% |
| Hormone Receptor | | |
| ER posritive | 76 | 86% |
| ER negative | 3 | 3% |
| PR posritive | 69 | 78% |
| PR negative | 10 | 11% |
| Her2 posritive | 3 | 3% |
| Her2 negative | 68 | 77% |
| Not Applicable | 9 | 10% |
| Menopausal Status | | |
| Pre-Menopausal | 19 | 22% |
| Peri-Menopausal | 7 | 8% |
| Post-Menopausal | 61 | 69% |
| Not Applicable | 1 | 1% |
